# Reducing methylation of histone 3.3 lysine 4 in the medial ganglionic eminence and hypothalamus recapitulates neurodevelopmental disorder phenotypes

**DOI:** 10.1101/2025.05.02.651761

**Authors:** Jianing Li, Anthony Tanzillo, Giusy Pizzirusso, Adam Caccavano, Ramesh Chittajallu, Mira Sohn, Daniel Abebe, Yajun Zhang, Ken Pelkey, Ryan K. Dale, Chris J. McBain, Timothy J. Petros

## Abstract

Methylation of lysine 4 on histone H3 (H3K4) is enriched on active promoters and enhancers and correlates with gene activation. Disruption of H3K4 methylation is associated with numerous neurodevelopmental diseases (NDDs) that display intellectual disability and abnormal body growth. Here, we perturb H3K4 methylation in the medial ganglionic eminence (MGE) and the hypothalamus, two brain regions associated with these disease phenotypes. These mutant mice have fewer forebrain interneurons, deficient network rhythmogenesis, and increased spontaneous seizures and seizure susceptibility. Mutant mice are significantly smaller than control littermates, but they eventually became obese due to striking changes in the genetic and cellular hypothalamus environment in these mice. Perturbation of H3K4 methylation in these cells produces deficits in numerous NDD-associated behaviors, with a bias for more severe phenotypes in female mice. Single cell sequencing reveals transcriptional changes in the embryonic and adult brain that underlie many of these phenotypes. In sum, our findings highlight the critical role of H3K4 methylation in regulating survival and cell-specific gene regulatory mechanisms in forebrain GABAergic and hypothalamic cells during neurodevelopment to control network excitability and body size homoeostasis.

## INTRODUCTION

Several decades of research have defined the critical role of chromatin remodelers and epigenetic modifying enzymes in regulating proper gene expression throughout development. There are ∼300 human genes coding for proteins that drive epigenetic modifications (readers, writers and erasers of epigenetic marks, chromatin remodelers, etc.), and these genes are highly intolerant to genetic variation^1^. Predictably, heterozygous loss-of-function variants in these genes leads to numerous neurodevelopmental diseases (NDDs)^2^, with many of these NDDs displaying a co-occurrence of intellectual disability (ID) and abnormal body growth^3,4^.

Many epigenetic machinery proteins target post-translational modifications of histones that are highly dynamic and required for proper transcriptional activation and repression^5,6^. One critical modification is methylation of histone H3 lysine 4 (H3K4), with trimethylation of H3K4 (H3K4me3) strongly correlating with promoter activation and gene expression. Dysregulation of H3K4me is associated with NDDs with ID, autism spectrum disorder (ASD), schizophrenia and substance-related disorders^7–9^. Canonical histones H3.1 and H3.2 are enriched in fetal mouse and human brains and decrease over time, with histone H3 variant H3.3 becoming the predominant histone H3 in postmitotic neurons that continues throughout adulthood^10–12^. H3.3 is encoded by 2 genes, *H3f3a* and *H3f3b*. Several studies have removed one or both genes in mice that lead to variable survivability and phenotypes depending on genetic targeting strategy, compensation and onset of gene loss^13^. Notably, conditional removal of both genes in the dorsal forebrain results in lethality several hours after birth^12^, complicating assessment of H3K4me function throughout neurodevelopment.

There are 6 members of the lysine-specific methyltransferase 2 (KMT2) family that methylate H3K4 in mammals (*KMT2A-D* & *F-G*), and 7 histone lysine demethylases that target H3K4me (*KDM1A-B*, *KDM2B* & *KDM5A-D*)^9,14^. Patients with variants in *KMT2A* (Wiedemann-Steiner syndrome), *KMT2C* (Kleefstra syndrome) and *KMT2D* (Kabuki syndrome) display ID, microcephaly, epilepsy and short stature^6,15–18^. Variants in demethylases *KDM5A-C* are associated with various forms of ID and ASD^9^, indicating a delicate balance H3K4 methylation is critical for normal brain development.

The comorbidities of epilepsy, ID and abnormal body growth in many of these H3K4me-associated diseases are indicative of altered excitatory-inhibitory balance and hypothalamic defects in these diseases. While several studies have explored decreased H3K4me in excitatory cells (mainly through loss of KMT2 genes)^19–23^, how disruption of H3K4me alters the fate and function of forebrain inhibitory interneurons and hypothalamic cells has not been explored. Interneurons are a highly diverse cell population that arise from two transient structures in the embryonic ventral forebrain, the medial and caudal ganglionic eminences (MGE and CGE, respectively)^24–26^. Abnormal development and function of interneurons has been linked to the pathobiology of NDDs such as schizophrenia, autism and epilepsy^27–30^, and many NDD-associated genes are enriched in interneuron progenitors^31–37^.

Here we devised a genetic strategy to reduce H3K4me specifically in the MGE and hypothalamus, two cell populations associated with common phenotypes of H3K4me dysregulation such as ID, epilepsy and control of body growth. Our results reveal significant genetic and cellular changes of MGE-derived interneurons and hypothalamic cells, leading to behavioral phenotypes associated with NDDs. In many assays, female mice display greater deficits compared to males. Electrophysiology recordings confirmed altered intrinsic properties of hippocampal interneurons and reduced network synchrony in mutant mice. In sum, our multimodal analyses reveal how a reduction of H3K4 methylation leads to both similar and distinct changes in forebrain interneurons and hypothalamic cells that, in part, underlie a variety of complex phenotypes related to NDDs.

## RESULTS

### Disruption of H3K4 methylation in the brain alters viability and body size

To manipulate H3K4me, we used a conditional transgenic mouse line *LSL-K4M* with a floxed-stop cassette followed by an HA-tagged human H3.3 sequence where the lysine 4 residue is mutated to a methionine (*H3.3K4M*)^38^, thus disrupting normal H3K4 methylation (Figure 1A). The transcription factor *Nkx2.1* is a ‘master regulator’ for all MGE-derived interneurons^39,40^ and is also expressed in the developing hypothalamus^41^ where it is critical for nuclei involved in regulating appetite and energy homeostasis^42–44^. We generated *Nkx2.1-Cre*;*H3.3K4M*;*Ai9* wild-type, heterozygous and homozygous mice (H3.3K4M WT, Het and Hom, respectively) to drive expression of H3.3K4M protein in the MGE and hypothalamus, with Nkx2.1-lineage cells expressing the red fluorescent reporter tdTomato. These H3.3K4M Hom mice contain both endogenous WT H3.3 genes (*h3f3a* and *h3f3b*) and the exogenous mutant H3.3K4M alleles, thus mimicking heterozygous disease-associated variants in humans that reduce but don’t eliminate H3K4 methylation (Figure 1B). We confirmed strong expression of HA-tagged H3.3K4M protein and a corresponding reduction of H3K4 methylation in the MGE and hypothalamus of *H3.3K4M* Hom embryos via immunohistochemistry and Western Blots (Figure 1C-E). H3.3 protein is strongly upregulated in postmitotic neurons and increases over time^10,12^ whereas *Nkx2.1* is expressed in cycling progenitors. Having H3.3K4M active prior to cell cycle exit is critical to substitute endogenous H3.3 with mutant H3.3K4M protein due to the prolonged half-life of histone proteins in the brain^45,46^. Additionally, this approach circumvents the compensatory problems that can arise from having 2 genes encoding H3.3 and 6 KMT2 family members that can methylate H3K4. H3.3K4M Hom mice were significantly smaller than WT mice during the first 6 postnatal weeks, with ∼25% of Hom mice dying during this period (Figure 2A-C). H3.3K4M Het mice were similar size as WT littermates, but a similar percentage of Het mice also died

**Figure 1.**
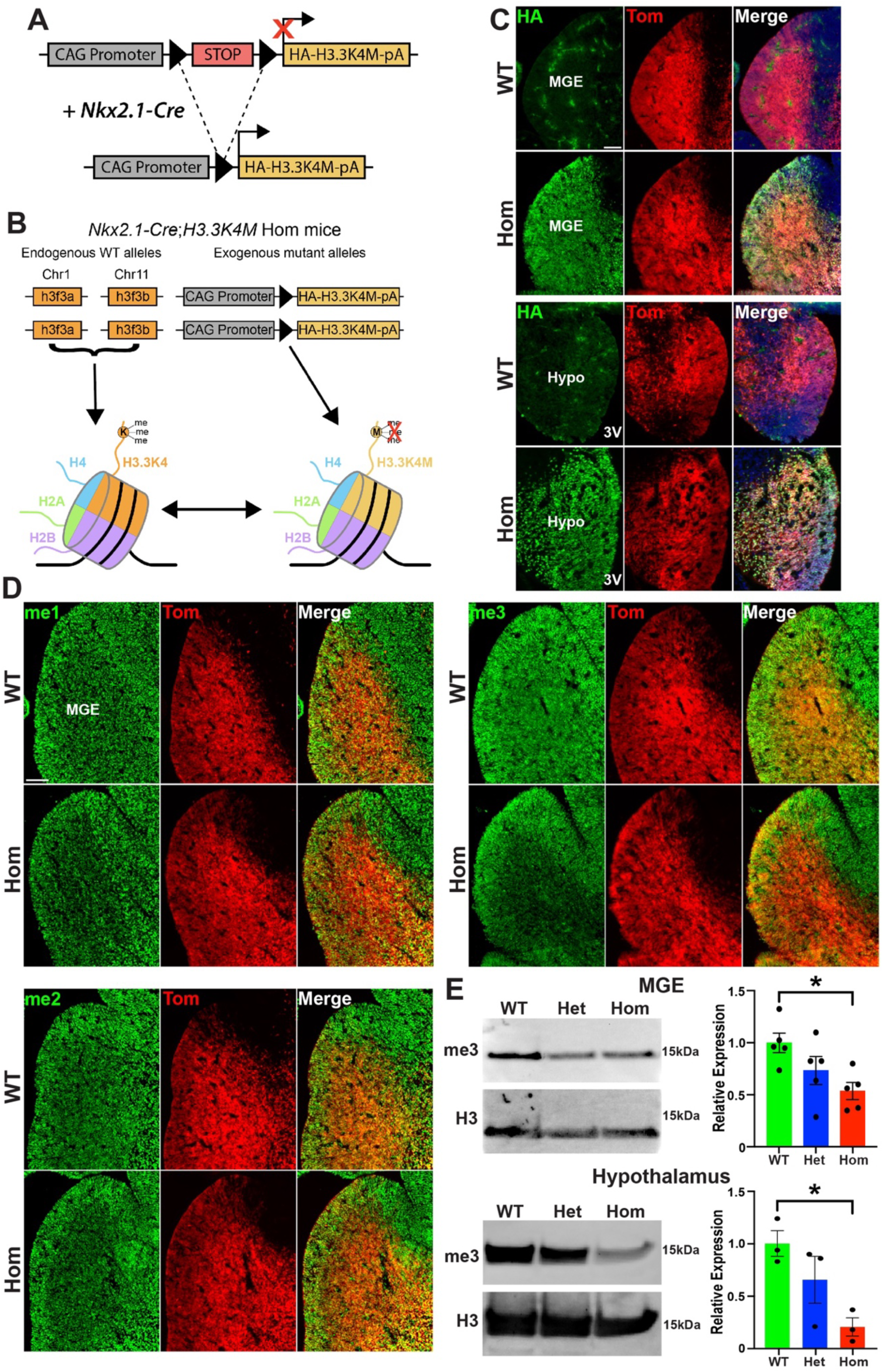
Downregulation of H3K4me3 in the H3.3K4M mouse. **A-B.** Schematic of *LSL-K4M* transgene (A), and combination with endogenous H3.3 genes leads to expression of both WT H3.3 and mutant H3.3K4M proteins (B). **C.** Upregulation of HA-tagged H3.3K4M protein in the MGE (top) and hypothalamus (bottom) of *Nkx2.1-Cre*;*H3.3K4M*;*Ai9* WT and Hom mice. 3V = 3^rd^ ventricle. **D.** Decrease of H3K4me1, H3K4me2 and H3K4me3 (me1, me2, me3, respectively) in the MGE of H3.3K4M Hom mice. **E.** Western blots of H3K4me3 levels in the MGE and hypothalamus with quantification. Scale bars = 100 µm. All stats are one-way ANOVA followed by Tukey’s multiple comparison tests: * = p ≤ .05.

**Figure 2.**
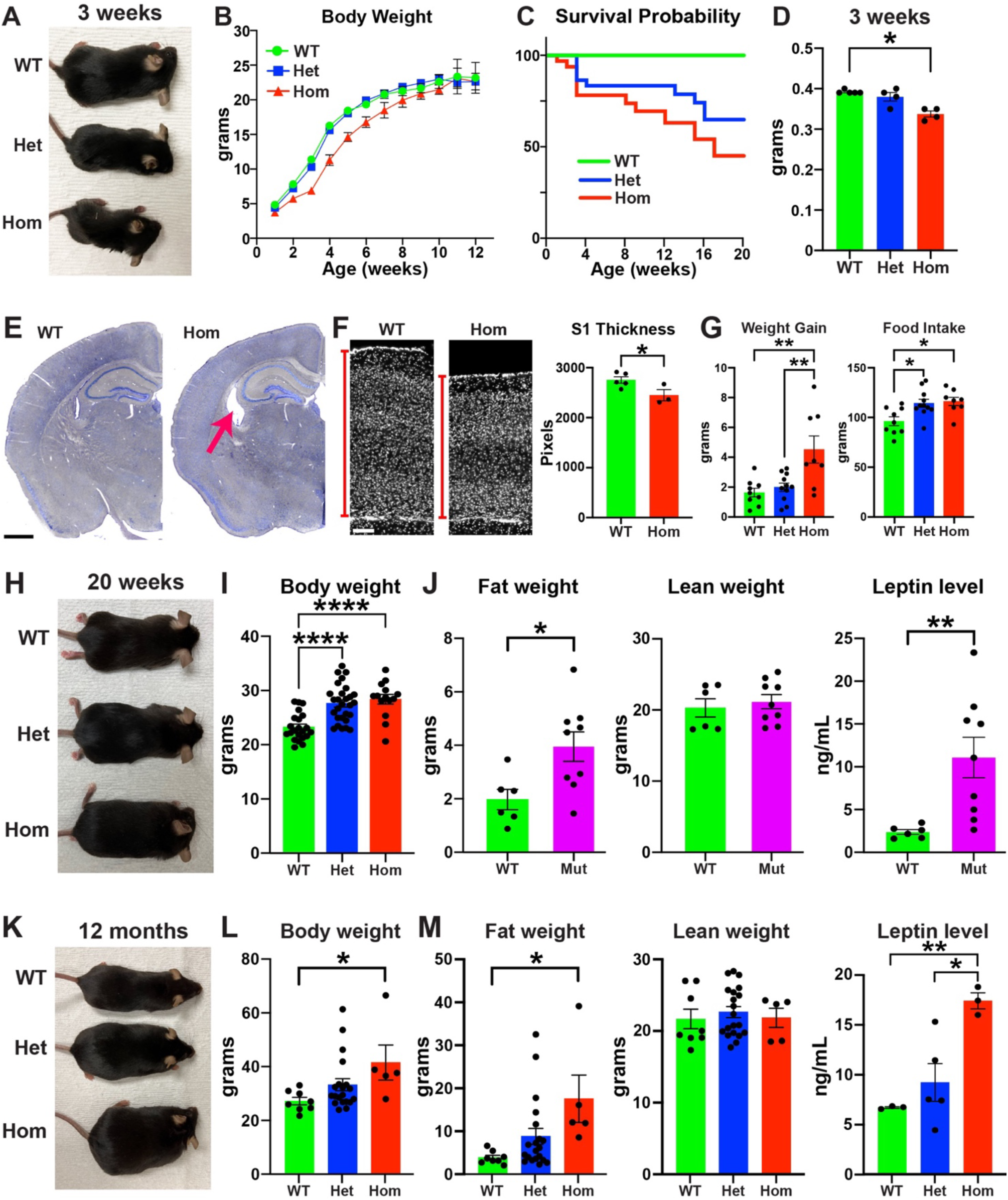
Disruption of H3K4 methylation alters viability and body size. **A-D.** Representative image of 3-week-old H3.3K4M WT, Het and Hom mice (A) with quantification of body weight (B), survival probability (C) and brain weight (D). **E.** Nissl staining reveals enlarged lateral ventricles (red arrow) in H3.3K4M Hom mice at 3 weeks. Scale bar = 1 mm. **F.** Representative DAPI image of somatosensory cortex (left) with decreased thickness in H3.3K4M Hom mice (right). Scale bar = 100 μm. **G.** Total weight gain (left) and food intake (right) from postnatal weeks 6-9. **H-I.** Representative image of 20-week-old H3.3K4M WT, Het and Hom mice (H) with quantification of body weight (I). **J.** Fat weight, lean weight and leptin levels of 20-week-old mice. **K-L.** Representative image of 12-month-old H3.3K4M WT, Het and Hom mice (K) with quantification of body weight (L). **M.** Fat weight, lean weight, and leptin levels of 12-month-old mice. Kruskal-Wallis followed by Dunn’s multiple comparisons (D) or one-way/two-way ANOVA followed by Tukey’s multiple comparison tests were performed based on normal distribution (B, G, I, L, M). Standard 2-tailed t-test (F, J: fat weight & lean weight) or Mann-Whitney test (J: leptin level) based on normal distribution when only WT and Hom mice tested: * = p ≤ .05, ** = p ≤ .005, **** = p ≤ .0001.

by 6 weeks. In addition to body weight, H3.3K4M Hom mice brains were also significantly smaller, had thinner cortices and had larger lateral ventricles compared to littermates (Figure 2D-F). While H3.3K4M Het and Hom mice continued to die over the next couple months (∼55% Hom and ∼30% Het die by 20 weeks), the survivors eventually surpassed the size of WT mice. From postnatal weeks 6-9, H3.3K4M Hom mice consumed more food and gained more weight compared to WT mice (Figure 2G). By 20 weeks, both H3.3K4M Het and Hom mice weighed more than WT mice with significantly higher fat weight and serum leptin levels (Figure 2H-J). This trend continued up to 1 year old mice where H3.3K4M Hom mice weighed ∼35% more, had > 4-fold more fat weight, and ∼2.5-fold higher serum leptin levels compared to WT (Figure 2K-M). Thus, perturbation of H3K4 methylation in the MGE and hypothalamus increases postnatal lethality and causes significant changes in brain size and body growth, likely due in part to increased food consumption and leptin levels.

### Decreased numbers and altered function of MGE-derived interneurons leading to increased seizure susceptibility

We observed that some H3.3K4M Hom mice had spontaneous seizures, suggesting a deficit in normal interneuron function. The MGE gives rise to nearly all parvalbumin- and somatostatin-expressing (PV+ and SST+, respectively) interneurons in the cortex, as well as a population of neuronal nitric oxide synthase-expressing (nNos+) neurogliaform and ivy cells in the hippocampus^24,25^. We observed a significant decrease in the density of Tom+ MGE-derived cortical interneurons in the cortex of P21 Het and Hom mice. This was primarily driven by a strong reduction in the density and proportion of PV+ interneurons, with a trend toward a reduction in SST+ interneuron density (Figure 3A-B). This decrease in PV+ cells was present in both the deep and superficial layers, implying a relatively homogeneous reduction throughout all cortical levels (Figure S1). Similar results were observed in the hippocampus, with a ∼40% reduction in MGE-derived interneurons affecting PV+, SST+ and nNos+ subtypes (Figure 3C-D). We also detected a population of Tom+/Olig2+ cells with small cell bodies in CA3, indicating that they are likely MGE-derived oligodendrocytes as previously described^47^. These Olig2+ oligodendrocytes were almost completely lost in the P21 H3.3K4M Hom mice, and we observed a corresponding decrease in myelin basic protein (MBP) in this region (Figure S2). Thus, disruption of H3K4 methylation in the MGE reduces interneurons in the cortex and hippocampus, with largest decreased in cortical PV+ and hippocampal nNos+ interneurons.

**Figure 3.**
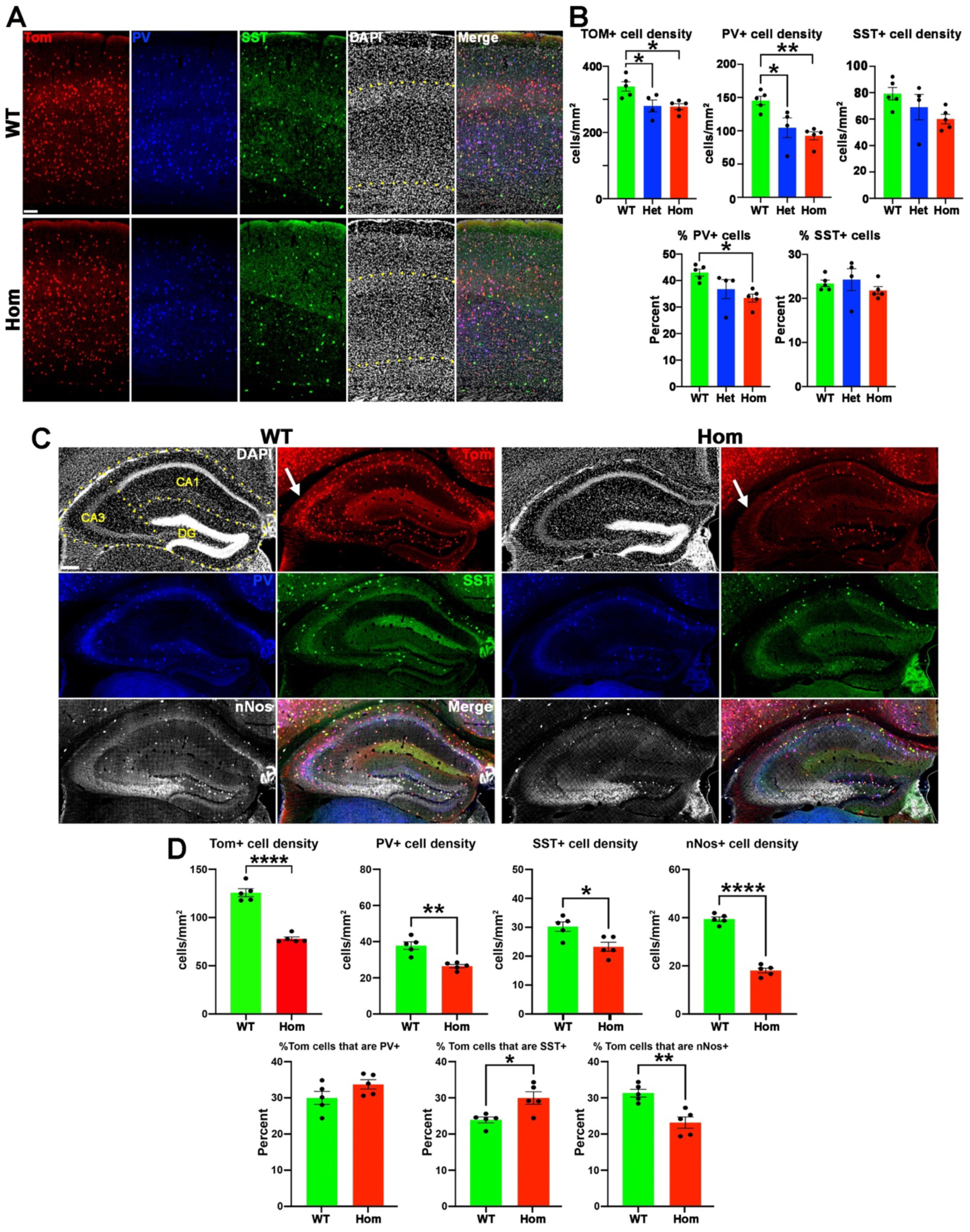
Decreased MGE-derived interneurons in H3.3K4M Hom mice. **A.** Representative images of P21 somatosensory cortex from *Nkx2.1-Cre*;*H3.3K4M*;*Ai9* WT and Hom mice stained for tdTomato (red), PV (blue), SST (green) and DAPI (white). Yellow dotted lines demarcate superficial and deep cortical layers. Scale bar = 50 μm. **B.** Quantification of cell density (top) and percent of PV+ and SST+ cells (bottom) in the somatosensory cortex. **C.** Representative images of P21 hippocampus in H3.3K4M WT and Hom mice stained for PV (blue), SST (green) and nNOS (white). Yellow dotted lines demarcate CA1, CA3 and dentate gyrus (DG) boundaries. Scale bar = 100 μm. **D.** Quantification of cell density (top) and percent of PV+, SST+ and nNos+ cells (bottom) in the hippocampus. All stats are one-way ANOVA followed by Tukey’s multiple comparison tests (B), or standard 2-tailed t-test when only WT and Hom mice tested (D): * = p ≤ .05, ** = p ≤ .005, *** = p ≤ .0005, **** = p ≤ .0001.

PV+ interneurons regulate feedforward inhibition that is critical for normal brain function and suppressing seizure activity^48,49^. Some P10-P14 H3.3K4M Hom mice displayed robust spasm-like behaviors (e.g., rapid, full flexions and extensions of limbs, trunk flexion, trunk curling and back arching) (Figure 4A & Supplementary Video 1). This epileptiform activity was also observed in 3-month-old H3.3K4M Hom mice, which exhibit head nodding, continuous whole-body myoclonus, rearing tonic seizure and tonic-clonic seizure with wild jumping (Figure 4A & Supplementary Video 2). This behavior was strongly sex-biased, with 70% of H3.3K4M Hom female mice displaying spontaneous seizures while only ∼14% of Hom males showed this behavior (Figure 4B). We did not observe any spontaneous seizure-like behavior in H3.3K4M Het mice.

**Figure 4.**
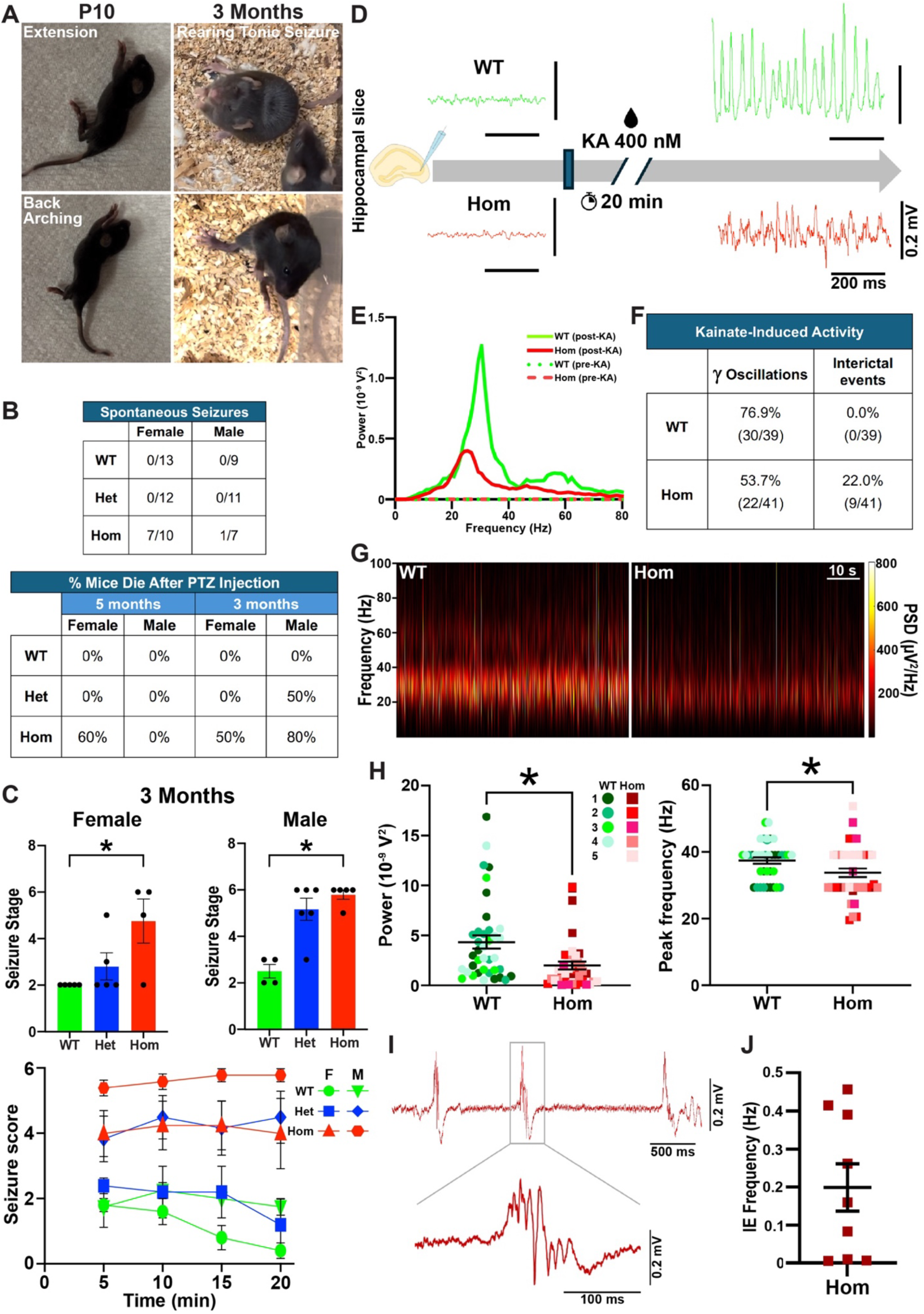
Spontaneous seizures and increased seizure sensitivity in H3.3K4M Hom mice. **A.** Still images depicting H3.3K4M Hom mice undergoing infantile spasms at P10 (left) and spontaneous seizures at 3 months (right). **B.** Quantification of spontaneous seizures observed in home cages from 3-5 months (top) and mortality rate after PTZ injection at 5 and 3 months (bottom). **C.** Maximum scores (seizure stage, top) for males (40 mg/kg) and females (20 mg/kg) following PTZ injection at 3 months and during the 20-minute observation period (bottom). **D-E**. Representative local field potential traces recorded in *ex vivo* hippocampal slices from WT (green) and H3.3K4M Hom (red) brains before (left) and after (right) KA exposure (D), with respective power spectra (E). **F.** Percent (and total number) of hippocampal slices displaying KA-induced gamma oscillations (left) and interictal events (right). **G.** Representative spectrograms of gamma oscillations from WT (left) and H3.34M Hom (right) brains. PSD: Power Spectra Density. **H.** Significant decrease in power (left) and peak frequency (right) in H3.3K4M Hom mice compared to WT. **I-J**. Representative trace showing interictal events (IE) (I), and IE frequency from 9 slices that displayed IE (J) in H3.3K4M Hom slices. Kruskal-Wallis followed by Dunn’s multiple comparisons (C: top) or two-way ANOVA followed by Tukey’s multiple comparison tests (C: bottom) were performed based on Normal distribution. Shapiro-wilk test was used to assess normality for electrophysiology data and unpaired T-test or Mann-Whitey test (H) followed accordingly: * = p ≤ .05.

To assess seizure susceptibility, we induced seizures with a single injection of Pentylenetetrazole (PTZ) at 5-months. We used a dose of 20 mg/kg, ∼50% of the typical effective concentration^50^, because some Hom mice already display spontaneous seizures. While both male and female 5-month-old H3.3K4M Hom mice displayed increased seizure susceptibility compared to WT, female Hom mice had more severe seizures, with 60% not recovering while no males died (Figure 4B & Figure S3A). Since Hom mice are already obese at 5-months, we also performed the PTZ injections at 3-months when WT and Hom mice are equal size. Since female mice are more sensitive to PTZ-induced seizures^51^, we increased the concentration of PTZ (40 mg/kg) for male mice. Both male and female Hom mice displayed increased seizure susceptibility at 3-months, with the increased dose in males generating similar results to female mice (Figure 4B-C).

The temporal lobe, and particularly the hippocampus, is the most common epileptic locus, and that is where we observe the greatest interneuron loss (Figure 3). To assess the function of MGE-derived interneurons, we performed whole cell patch-clamp recordings on Tom+ cells in *ex vivo* hippocampal slices from 8-week-old female WT and H3.3K4M Hom mice. Quantifying six intrinsic properties (input resistance, Tau, capacitance, Sag ratio, maximum firing rate, AP half-width) was sufficient to define the 3 expected interneuron cell types via unbiased hierarchical clustering and K-means clustering (Figure S3B-D). Consistent with our immunohistochemical findings, putative PV+/fast-spiking (FS), SST+/non-FS (NFS), and nNos+/slow-spiking (SS) interneurons were observed in H3.3K4M Hom mice with properties largely similar to those in WT mice. However, at the population level, all three interneuron subtypes in H3.3K4M Hom mice showed greater variability in intrinsic properties compared to WT, leading to less refined clustering on the PCA maps (Figure S3D). This increased variance was statistically significant in some instances, such as the firing rate and capacitance in FS cells (Figure S3E). Such increased variability in electrical properties is reminiscent of the developing hippocampus^52,53^, suggesting interneuron maturation deficits in H3.3K4M Hom mice.

Cortical circuit information coding requires precision in the timing, extent and synchrony of activity within glutamatergic principal cell assemblies that is largely orchestrated by local circuit interneurons. Brain oscillations in the gamma-frequency band are critical for higher cognitive function and critically rely on balanced phasic excitatory and inhibitory drive^54–58^. To examine if the interneuron deficits observed in H3.3K4M Hom mice disrupt network rhythmogenesis, we probed kainate (KA)-induced gamma oscillations in *ex vivo* hippocampal slices from 3-month-old mice male and female mice (Figure 4D-E). Gamma oscillations were detected in 77% of slices from WT mice but only 54% of slices from H3.3K4M Hom mice (Figure 4F), suggesting a reduced propensity for physiological network rhythmogenesis in H3.3K4M Hom mice. Slices from H3.3K4M Hom mice displayed significantly decreased gamma power and peak frequency compared to WT mice (Figure 4E-H), indicating that H3.3K4M slices produce smaller and slower oscillations. Additionally, multiple slices from H3.3K4M mice developed KA-induced epileptiform bursts, which were not observed in any slices from WT mice (Figure 4F, I-J).

This finding suggests unrestrained network hyperexcitability in H3.3KM Hom mice, consistent with the increased seizure susceptibility observed *in vivo* (Figure 4C & S3A). As widespread perisomatic inhibition is critical for both circuit gamma entrainment and seizure control^48,59^, it is likely that the deficits in hippocampal FS/PV+ cells in H3.3K4M mice directly contribute to disrupted rhythmogenesis and epilepsy. Therefore, perturbation of H3K4 methylation decreases the number of MGE-derived interneurons, alters their intrinsic properties and disrupts normal network activity, leading to spontaneous seizures and increased seizure susceptibility; this is similar to temporal lobe epilepsy (TLE) patients and animal models of TLE^60–63^.

### Increased anxiety and impaired locomotor activity in H3.3K4M Hom mice

We performed a series of behavior tests on either juvenile (5-11 weeks) or adult (3-5 months) H3.3K4M WT and mutant mice (Figure S4A) to determine if disruption of H3K4 methylation in the MGE and hypothalamus induces behavioral deficits associated with NDDs. To assess anxiety behaviors, we characterized mice in the open field and light-dark box tests. Both female and male H3.3K4M Hom mice spent significantly less time in the center zone of the open field compared WT mice (Figure 5A-B). Female Hom mice displayed a trend of increased average speed in the open field test, which was not observed in male mice (Figure 5B). In the light-dark box test, H3.3K4M Hom male and female male mice spent more time in dark zone compared to WT mice (Figure 5C). These two tests confirmed increased anxiety-like behavior with reduced H3K4 methylation.

**Figure 5.**
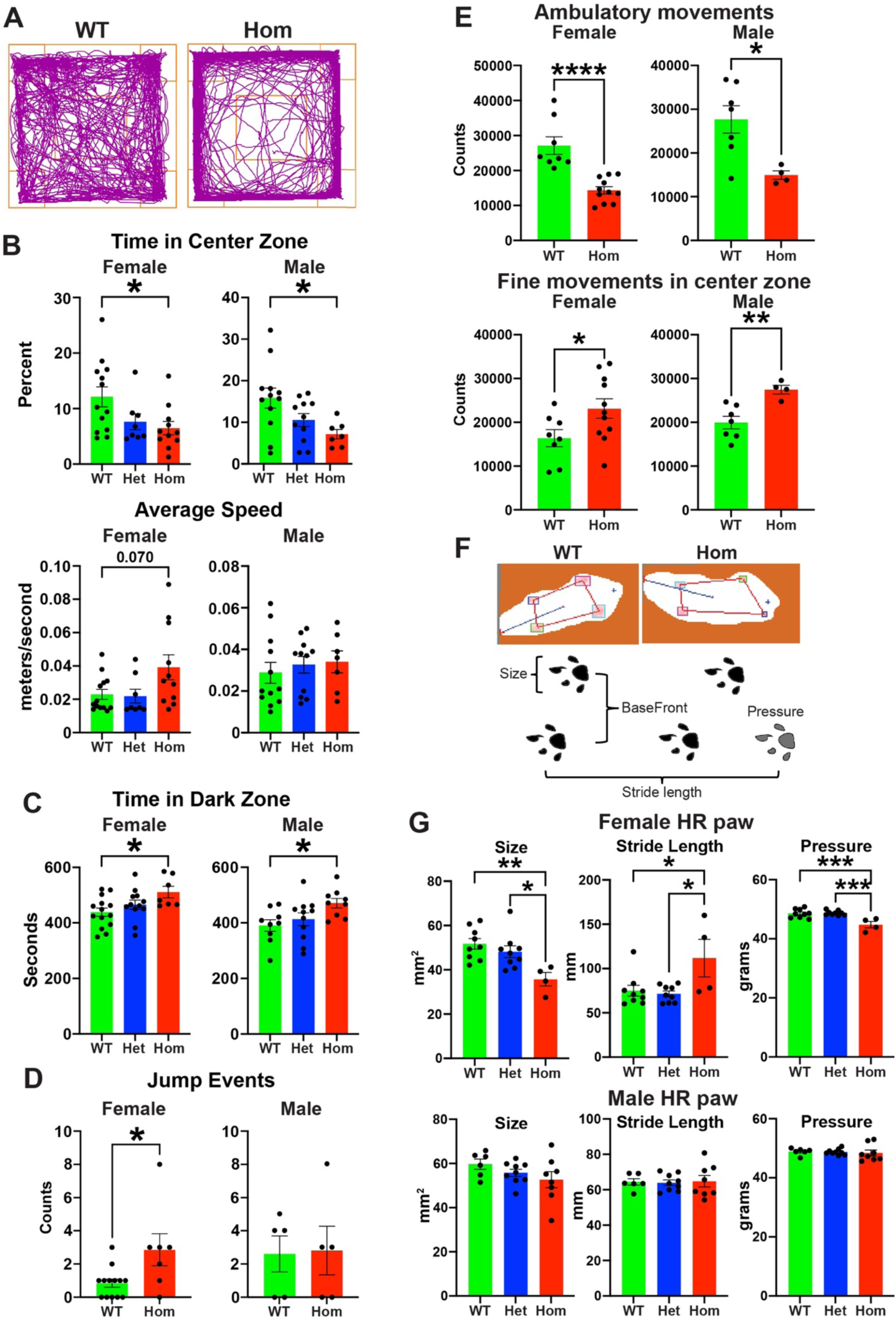
H3.3K4M Hom mice display increased anxiety and impaired locomotion. **A.** Representative trajectory plots of H3.3K4M WT and Hom mice in the open field test. Center zone indicated by orange square. **B.** Time in the center zone (top) and average speed (bottom) in the open field test. **C.** Time spent in the dark zone of the light-dark box. **D.** Jump events recorded in the Cliff Avoidance Reaction (CAR) test. **E.** Beam break counts of ambulatory movements in whole zone (top) and fine movements in the center zone (bottom) in home cage over 4 days. **F.** Representative images from free walk test (top) with schematic depicting gait measurements (bottom). **G.** Size, stride length and pressure of hind right (HR) paw of in the free walk test. All stats are one-way ANOVA followed by Tukey’s multiple comparison tests when WT, Het and Hom mice (B, C, G). Standard 2-tailed t-test when only WT and Hom mice tested (D, E): * = p ≤ .05, ** = p ≤ .005, *** = p ≤ .0005, **** = p ≤ .0001.

Since H3.3K4M Hom mice were observed jumping out of the cage during normal handling, we assessed these mice for increased impulsivity. Impulsivity is a primary clinical symptom of ADHD and is observed in other NDDs^64,65^. We used the cliff avoidance reaction (CAR) test to assess impulsivity in these mice. Female H3.3K4M Hom mice had significantly more jump off events from the platform compared with WT (Figure 5D & S4B), suggesting increased impulsive-like behaviors. There was no difference in male mice.

We observed that some H3.3K4M Hom mice displayed unsteady gait and altered locomotion. To assess general locomotor activity, we continuously recorded singly housed mice for 4 days using the Photobeam Activity System. Both H3.3K4M Hom female and male mice displayed decreased ambulatory movements compared to WT mice, with an increase in fine movements in the center zone (Figure 5E & Figure S4C). No difference in rearing activity was observed (Figure S4C). We used the Free-Walk system to quantify gait parameters and motor coordination in 3-month-old mice (Figure 5F). Female H3.3K4M Hom mice displayed significantly reduced size, pressure, stance of hind-right paw, decreased BaseFront, and an increased stride length compared to WT mice (Figure 5G & Figure S4D). Gait properties of male H3.3K4M Hom mice were normal, except for a slight increase in BaseFront. Thus, H3.3K4M Hom mice display increased anxiety, decreased general locomotion and altered gait activity, with females generally displaying more severe symptoms compared to males.

### H3.3K4M Hom mice display impaired sensorimotor gating and memory

We then assessed higher order cognition and behavior in H3.3K4M mice. Proper sensorimotor gating is a critical aspect of normal brain function and animal behavior, and deficits in prepulse inhibition (PPI) are observed in NDDs such as schizophrenia and obsessive-compulsive disorder^66^. In the acoustic startle reflex test, female H3.3K4M Hom mice showed an increase in startle amplitude and a decrease percent of PPI in a series of prepulse intensities compared to WT mice (Figure 6A). No significant differences were observed in the male H3.3K4M Hom mice.

**Figure 6.**
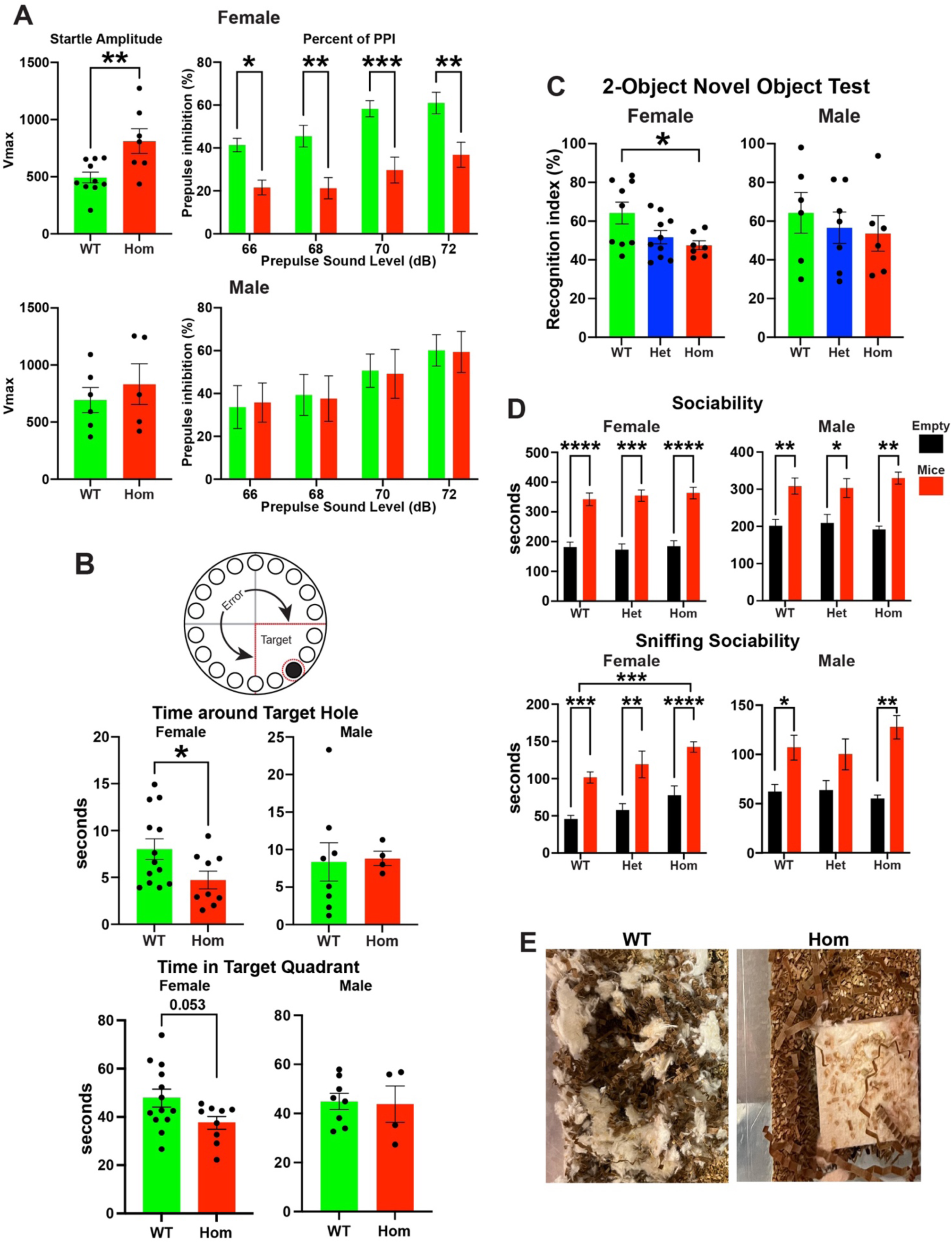
H3.3K4M Hom mice display impaired sensorimotor gating and memory. **A.** Startle amplitude (left) and percent prepulse inhibition (PPI) (right) in H3.3K4M WT and Hom mice. **B.** Barnes maze schematic showing error and target zones (top), with red dotted line indicates target quadrant and region around target hole used for quantification. Time spent around target hole (middle) and in target quadrant (bottom) during the probe test. **C.** Recognition index in the 2-object novel object recognition test. **D.** Time spent in the chambers with (red) or without (black) stimulus mouse in the three-chamber test (top). Time spent sniffing at the wire cage with or without stimulus mouse (bottom). **E.** Representative images of nest building for singly housed male H3.3K4M WT and Hom mice. All stats are one-way ANOVA (C) or two-way ANOVA (A: percent of PPI, D) followed by Tukey’s multiple comparison tests when WT, Het and Hom mice. Standard 2-tailed t-test when only WT and Hom mice tested (A: startle amplitude, B): * = p ≤ .05, ** = p ≤ .005, *** = p ≤ .0005, **** = p ≤ .0001.

We utilized the Barnes maze and 2-object novel recognition paradigms to explore memory deficits in H3.3K4M mice. In the Barnes maze, mice are challenged to find and remember the location of an escape hole in an elevated maze while motivated by loud white noise. Female H3.3K4M Hom mice exhibited a significantly shorter time around the target hole and trended to spend less time in the target quadrant compared to WT mice (Figure 6B), indicating decreased memory performance in this task. Notably, H3.3K4M Hom mice traveled a greater total distance compared to WT mice, indicative of hyperactivity and increased anxiety (Figure S4E). In the 2-object novel recognition assay, WT female mice exhibited preference for novel versus familiar objects as expected, while female H3.3K4M Hom mice showed no preference for the novel object (Figure 6C). No significant differences were observed in the male H3.3K4M Hom mice.

To evaluate sociability deficits in these mutant mice, we performed the three-chamber test. We found no significant differences in sociability between H3.3K4M Hom and WT mice (Figure 6D & Figure S4F). Curiously, H3.3K4M Hom female mice spent more time directly interacting with (sniffing) both the empty object and mice compared WT. Last, we observed nest building defects in most singly housed H3.3K4M female and male Hom mice (Figure 6E), which is observed in numerous mouse models of NDDs^67,68^.

Altogether, our behavioral data indicate that perturbation of H3K4 methylation in the MGE and hypothalamus produces a broad range of behavioral phenotypes, many of which are commonly found in NDDs and other models of altered H3K4 methylation^8,9^. H3.3K4M Hom mice display increased anxiety and impulsivity, hypoactivity and decreased ambulatory locomotion in home cage, abnormal gait, and deficits in sensorimotor gating and memory. Notably, female Hom mice displayed more severe phenotypes compared to males in many assays, indicating a clear sex bias in these mutant mice.

### Alterations in transcriptome associated with interneurons fate and seizure

To define genetic and cellular changes arising from disruption of H3K4 methylation that underlie the observed phenotypes, we performed single nucleus Multiome analysis (snRNA-seq + snATAC-seq) on E13.5 MGE, E13.5 hypothalamus, P60 cortical MGE-derived interneurons and P60 hypothalamus cells from male and female WT and H3.3K4M Hom mice. For these experiments, we generated *Nkx2.1-Cre*;*H3.3K4M*;*Sun1-sfGFP* mice, allowing us to harvest *Nkx2.1*-lineage GFP+ nuclei via flow cytometry. After QC filtering, we obtained a total of 50,684 nuclei from embryonic mice and 22,116 nuclei from P60 mouse brains (Supplementary Table 1). There was clear segregation of both ages and brain regions when visualizing RNA-only, ATAC-only or integrated RNA and ATAC data using the weighted nearest neighbor (WNN) analysis (Figure 7A and Figure S5A-D). Cells from the same age and brain region largely overlapped regardless of genotype or sex (Figure 7B and Figure S5E-F), indicating gross similarities in cell types between genotypes and sexes.

**Figure 7.**
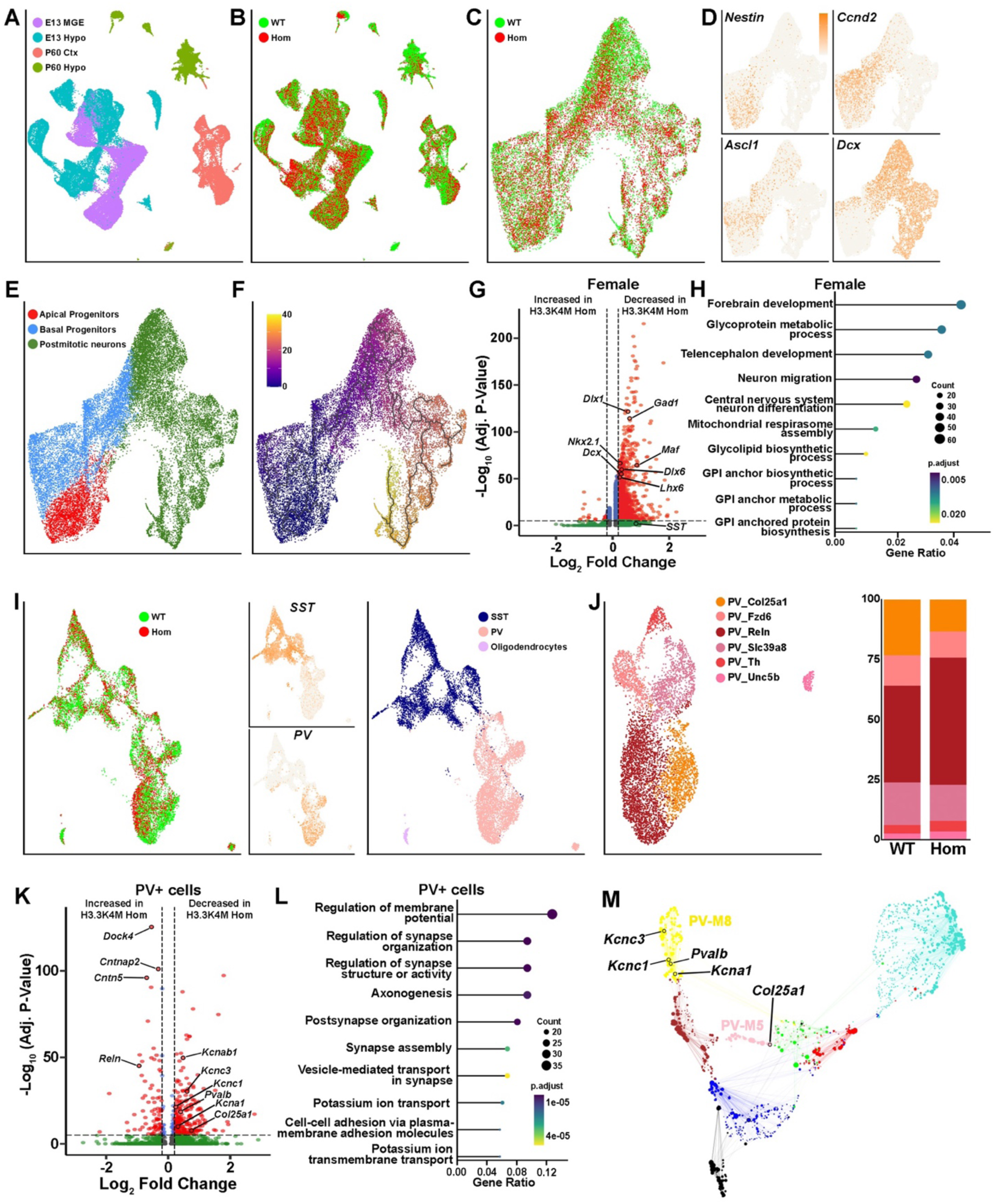
Altered transcriptomes of interneurons associated with neuronal maturation and epilepsy. **A-B.** UMAP plots of integrated single nuclei RNA and ATAC data via weighted nearest neighbor (WNN) of Nkx2.1-lineage cells from E13.5 MGE, E13.5 hypothalamus, P60 cortex and P60 hypothalamus annotated by age and tissue (A), and genotypes (B). **C-F.** Integrated RNA and ATAC UMAP plots of E13.5 MGE cells annotated by genotypes (C), marker genes for radial glia cells/apical progenitors (*Nestin*), basal progenitors (*Ccnd2* and *Ascl1*) and post-mitotic immature neurons (*Dcx*) (D), clusters defined as apical progenitors, basal progenitors and postmitotic neurons (E), and pseudotime developmental trajectory (F). **G.** Volcano plot depicting genes significantly increased or decreased in E13.5 H3.3K4M Hom female MGE compared to WT. **H.** clusterProfiler GO enrichment top hits of biological process for DEGs of E13.5 female MGE. **I.** UMAP plots of Nkx2.1-lineage, MGE-derived cells in the P60 cortex cells annotated by genotypes (left), *SST* and *PV* genes (middle) and SST+ and PV+ clusters (right). **J.** PV+ cluster replotted and annotated by 6 PV+ subtypes (left), and relative proportions of PV+ subtypes in H3.3K4M WT and Hom cortices (right). **K.** Volcano plot depicting DEGs in P60 PV+ interneurons between H3.3K4M WT and Hom mice. **L.** clusterProfiler GO enrichment top items of biological process for DEGs of P60 PV+ interneurons. **M.** Network visualization of the 8 PV modules identified by WGCNA, with several hub genes labeled.

In the E13.5 MGE cells, we identified 20 putative cell clusters (Figure 7C and Figure S6A). Cells could be classified as apical progenitors (*Nestin*+), basal progenitors (*Ccnd2*+ and *Nestin-*) and postmitotic neurons (*Dcx*+ and *Ccnd2-*) that followed the expected developmental trajectory (Figure 7D-F and Figure S6B). We consistently observed a smaller percentage of GFP+ nuclei in the MGE from H3.3K4M Hom mice (75.3% in WT vs. 62.1% in Hom), which is consistent with fewer interneurons in the cortex and hippocampus in mutant mice. We performed differential gene expression and differential peak accessibility analysis to identify how perturbation of H3K4 methylation alters gene expression. We identified many differentially expressed genes (DEGs) in both H3.3K4M Hom female and male MGE, with the overwhelming majority of DEGs being downregulated in Hom mice (Figure 7G and Figure S6C). Nearly all differentially accessible peaks were also less accessible in the MGE of H3.3K4M Hom mice (Figure S5G). Many genes critical for general interneuron development (*Dlx1*, *Dlx6, Gad1)* and specific for MGE-derived interneurons (*Nkx2.1, Lhx6*, *Lhx8*) (Figure 7G and S6C) were downregulated in Hom mice. Of note, 2 genes predictive of PV+ interneurons, *Maf* and *Mef2c*^69,70^, are reduced in H3.3K4M Hom MGE while SST is unchanged, which is consistent with the specific reduction of PV+ cells in the adult cortex (Figure 3). Gene ontology (GO) analysis revealed that DEGs were enriched in categories relating to forebrain development and neuron migration (Figure 7H and Figure S6D).

To analyze transcriptional changes in mature MGE-derived interneurons in the adult, we subsetted the P60 cortical cells. These cells were cleanly divided into SST+ and PV+ interneurons, with a small population of oligodendrocytes (Figure 7I and Figure S6E). We focused on PV+ interneurons since they showed the greatest changes in H3.3K4M Hom mice in terms of cell numbers (Figure 3) and function (Figure 4). Based on previous PV+ classifications^71^, we identified six putative PV+ interneuron subtypes and noted a shift in the relative proportion of specific subtypes in H3.3K4M Hom mice: an increase of the PV-Reln population (from 40.3% to 53.0%) and a concomitant reduction in PV-Col25a1 cells (from 23.3% to 13.4%) (Figure 7J & Figure S6F). We confirmed this predicted increase of PV+/Reln+ cells in the cortex of H3.3K4M Hom mice (Figure S7). Genes related to cell-cell adhesion (*Dock4, Cntnap2, Cntn5*) were strongly upregulated in PV+ cells in H3.3K4M Hom mice, and the top gene families were related to cell adhesion and synapse formation (Figure 7K-L). Several genes involved in potassium ion transport (*Kcnab1, Kcna1, Kcnc1, Kcnc3*) were downregulated in the mutant, as was *Pvalb* (Figure 7K-L & Figure S6G). Of note, *Reln* was upregulated in the mutant whereas *Col25a1* was downregulated, which correlates with the proportional change in these PV+ subtypes (Figure 7K). Comparing PV+ cells from male and female H3.3K4M Hom mice revealed that many potassium ion transport associated genes (e.g., *Kcnh7, Kcnmb2, Kcnj3*) were decreased in females (Figure S6H), which is consistent with female mice being more susceptible to seizures (Figure 4B & Figure S3A).

To explore changes in gene regulatory networks upon perturbation of H3K4 methylation, we performed weighted gene co-expression network analysis (WGCNA)^72,73^ on PV+ interneurons. Eight co-expression modules were found in PV+ interneurons, with 5 of these modules being differentially regulated between WT and H3.3K4M Hom mice (Figure S6I). Notably, the downregulated potassium transporters genes *Kcna1, Kcnc1* and *Kcnc3* were found in module PV-M8, which has the largest change between WT and H3.3K4M Hom mice (Figure 7M). *Col25a1* was found in another module downregulated in Hom mice, PV-M5, which matches the reduction of PV-Col25a1 subtype. Potassium ion channels, and particularly Kv3 channels, are highly expressed in PV+ interneurons, and loss of Kv3 channels alters their fast-spiking properties and increases seizure susceptibility^74,75^, which matches the behavioral and electrophysiological changes in H3.3K4M Hom mice.

We observed fewer DEGs in SST+ interneurons compared to PV+ (208 in SST+ vs. 336 in PV+) (Figure S6J). Genes related to neuron cell adhesion and regulation of synapse organization (*Lama4*, *Cdh8*, *Nlgn1*, *Sparcl1* and *Epha4*) were downregulated in SST+ cells from H3.3K4M Hom mice, which could indicate altered synaptic connectivity with pyramidal neurons.

In sum, perturbation of H3K4 methylation alters the transcriptome of embryonic MGE and adult MGE-derived interneurons that are highly correlative, and likely causative, to interneuron-related phenotypes in the mutant mice. This includes decrease of PV-indicative genes in the MGE, changes in the proportion of PV+ subtypes, and downregulation of potassium ion transport, channels and gene modules that are associated with seizure susceptibility.

### Dysregulation of hypothalamic gene expression, cellular composition and structure in H3.3K4M mutant mice

The hypothalamus is the primary brain region regulating metabolism, feeding, stress response and other functions altered in H3.3K4M Hom mice. We subsetted the E13.5 hypothalamus datasets and used gene expression studies^76–78^ to identify *Nes*+ neural progenitor cells (NPCs) and 11 putative hypothalamic nuclei consisting of both glutamatergic (*Slc17A6*+) and GABAergic (*Slc32a1*+) cells (Figure 8A-C & Figure S8A-C). Similar to the MGE, we observed a lower percentage of GFP+ nuclei in the H3.3K4M Hom hypothalamus compared to WT during sorting (41.5% GFP+ nuclei in WT vs 25.5% in Hom). There was a more even distribution of upregulated and downregulated DEGs compared to MGE, with a notable increase in differentially accessible peaks in the H3.3K4M Hom mouse (Figure 8D & S5G). We observed an increased proportion of NPCs in H3.3K4M Hom mice (27.1% vs. 16.3%) (Figure 8C). Numerous genes associated with mitotic cell cycle phase transition (*Ccnd2, Ccnd1, Cdk6, Mcm6*) were upregulated in the H3.3K4M Hom hypothalamus (Figure 8D-E). Additionally, there is increased accessibility at the *Nes* promoter and intron enhancer in the hypothalamus of H3.3K4M Hom mice compared to WT (Figure 8F), suggesting that decreased H3K4me increases the proportion of *Nes*+ NPCs resulting in reduced or delayed neurogenesis.

**Figure 8.**
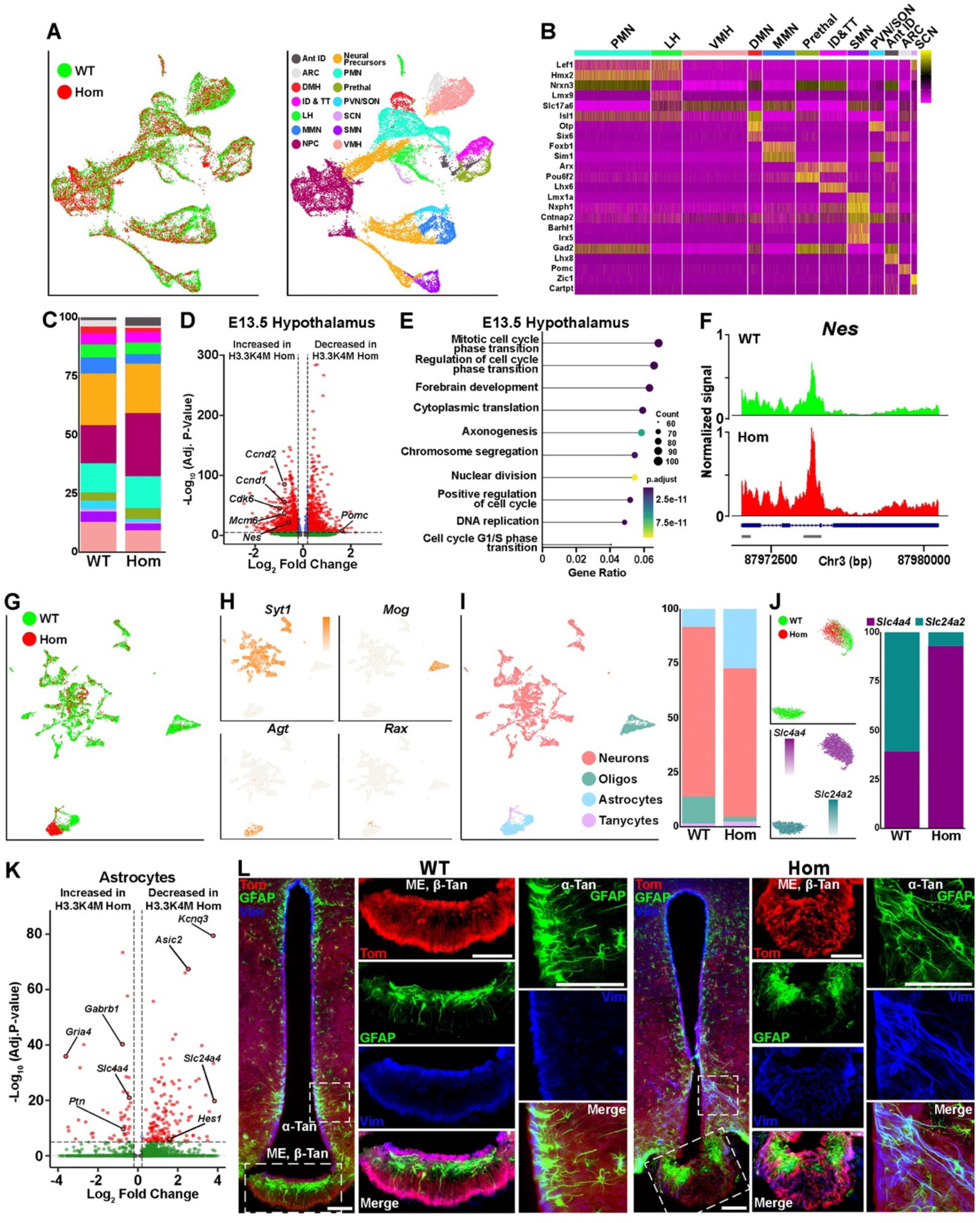
Significant changes in gene expression and cellular composition in the hypothalamus of H3.3K4M Hom mice. **A.** UMAP plots of integrated RNA and ATAC data of Nkx2.1-lineage cells from E13.5 hypothalamus annotated by genotypes (left) and putative hypothalamus nuclei (right). **B.** Heatmap depicting genes enriched in specific hypothalamic nuclei. **C.** Relative proportions of hypothalamic nuclei in H3.3K4M WT and Hom. **D.** Volcano plot depicting DEGs in E13.5 hypothalamus between H3.3K4M WT and Hom. **E.** clusterProfiler GO enrichment top biological process for DEGs of E13.5 hypothalamus between H3.3K4M WT and Hom. **F.** Genomic tracks showing increased peak at *Nes* promoter and intronic enhancer (marked by gray bars) in E13.5 H3.3K4M Hom hypothalamus. **G-H.** Integrated RNA and ATAC UMAP plots of P60 hypothalamus annotated by genotypes (G), and marker genes for neurons (*Syt1*), oligodendrocytes (*Mog*), astrocytes (*Agt*) and tanycytes (*Rax*) (H). **I.** UMAP plot annotated by cell type (left) and proportion of each cell type in WT and Hom mice (right). **J.** UMAP plots of astrocytes annotated by genotype (top) and genes demarcating these two distinct populations, *Slc4a4* and *Slc4a2* (bottom). Relative proportion of WT and Hom cells in each cluster (right). **K.** Volcano plot depicting DEGs in P60 astrocytes population between H3.3K4M WT and Hom. **L.** Representative images of ∼P60 hypothalamus from *Nkx2.1-Cre*;*H3.3K4M;Ai9* WT and Hom mice stained for GFAP (green) and Vimentin (blue). White boxes indicate higher magnification areas of median eminence (ME) and hypothalamus lining 3^rd^ ventricle for each genotype. Scale bars = 100 μm. Standard 2-tailed t-test was tested: * = p ≤ .05. PMN, premammillary nucleus; VMH, ventromedial hypothalamus; LH, lateral hypothalamus; DMN, dorsomedial nucleus; Prethal, Prethalamus; ARC, Arcuate nucleus; ID&TT, intrahypothalamic diagonal & tuberomammillary terminal; SMN, supramammillary nucleus; PVN/SON, paraventricular nucleus/supraoptic nucleus; Ant ID, Anterior intrahypothalamic diagonal; SCN, suprachiasmatic nucleus; MMN, mammillary nucleus.

Conversely, nuclei with the greatest proportional reduction in the H3.3K4M Hom mice were the arcuate nuclei (ARC), which is critical for regulating energy metabolism and food intake^79^, and the paraventricular nucleus/supraoptic nucleus (PVN/SON) nuclei that are involved in autonomic responses involving stress, body growth and hormone production^80^ (Figure 8C and Figure S8D). The ARC marker gene *Pomc* was decreased in the Hom hypothalamus (Figure 8D), consistent with a decrease proportion of ARC-fated nuclei.

In the P60 hypothalamus, we identified 40 putative cell clusters with clear separation of neurons (*Syt1*), astrocytes (*Agt*), oligodendrocytes (*Mog*) and tanycytes (*Rax*) (Figure 8G-I and Figure S8E-F). Specific hypothalamic nuclei could not be cleanly resolved in our dataset because we obtained too few cells relative to the heterogeneity of cell types and nuclei in the adult hypothalamus (Supplementary Table 1); a recent study compiled ∼400,000 mouse hypothalamic cells and identified 185 distinct cell clusters^81^. We observed a small number of DEGs in adult hypothalamic neurons, with the majority being increased in H3.3K4M Hom mice (Figure S8G). While genes labeling specific populations of ARC neurons (*Pomc*, *Agrp*, *Npy*) were not significantly changed, several genes associated with NADH or glycolysis metabolic processed (*Gapdh*, *Pfkp, Tpi1*) were increased in H3.3K4M Hom adult neuron (Figure S8G-H).

In the H3.3K4M Hom mice, we found a striking increase in the proportion of astrocytes (8.2% in WT vs. 27.3% in Hom) and decrease in oligodendrocytes (12.3% in WT vs. 2.2% in Hom) (Figure 8I). We confirmed an increase in hypothalamic GFAP+/Tom+ Nkx2.1-lineage astrocytes, and overall astrocyte numbers, in both male and female H3.3K4M Hom mice (Figure S9A-C). This increase in astrocytes is also found in diet-induced obese mice^82^. Unlike other cell types, hypothalamic astrocytes from WT and Hom mice form distinct clusters in the UMAP plots (Figure 8G). Re-clustering the astrocyte population revealed 2 distinct groups: a WT-only *Slc24a2*+ population and a mixed *Slc4a4*+ population (Figures 8J & S8I). Astrocytes play critical roles in controlling glucose metabolism and energy balance^44,83,84^, maintaining homeostasis of neurotransmitter and ions^85^, and regulating synapse plasticity linked to behaviors^86^. DEGs in astrocytes includes those involved in regulation of monoatomic ion transport (*Asic2*, *Gria4, Gabrb1*), glial cell differentiation (*Hes1*, *Ptn*) and regulation of membrane potential (*Kcnq3*, *Slc4a4* and *Slc24a4*), which may indict disruption to normal hypothalamic homeostasis in H3.3K4M Hom mice (Figures 8K & Figure S8J).

In sum, perturbation of H3K4 methylation alters the transcriptome of embryonic and adult hypothalamic cells, with a notable increase in the proportion of NPCs and reduction in the proportion cells in ARC and PVN/SON nuclei, which play critical roles in feeding behavior and growth hormone regulation. We also observed a large increase in hypothalamic astrocytes in H3.3K4M mice and distinct transcriptional astrocyte subtypes that likely alter hypothalamic function.

### Abnormal cellular organization in the hypothalamus of H3.3K4M Hom mice

Tanycytes are specialized glial cells within the blood-brain barrier of the hypothalamus that can detect nutrients and metabolites in the blood and cerebrospinal fluid, sending this information to the ARC and VMH to regulate body weight and energy metabolism^87,88^. They also act as neural progenitor cells, giving rise to new neurons and glia cells in the adult hypothalamus^87^. In WT mice, Nkx2.1-lineage Tom+/Vimentin+ (Vim+) tanycytes line the third ventricle (α-tan) and median eminence (ME, β-tan), with their processes extending out in an organized manner as previously described^89^ (Figure 8L). A layer of Tom-/GFAP+ astrocytes lie just below these tanycytes in the ME. In male and female H3.3K4M Hom mice, this organization is completely disrupted, with Vim+ tanycytes scattered throughout the ME and GFAP+ astrocytes clumped at the lateral edges of the ME (Figure 8L). α-tan processes along the wall of the third ventricle are more disorganized and fasciculated compared to WT mice. Additionally, myelin from oligodendrocytes normally lines the dorsal portion of the ME where it is involved in the regulation of energy availability and leptin sensitivity^90^. This myelin band was present in the ME of WT mice but was absent in Hom mice, with myelin instead restricted to the lateral edges of the ME, similar to GFAP+ astrocytes (Figure S9D).

We subsetted out the *Rax*+/*Col23a1*+ tanycytes in the hypothalamus and identified the 2 tanycyte subtypes as previously described^91^: *Vcan*+ α-tans and *Col25a1*+ β-tans (Figure S8K). We identified very few DEGs in tanycytes, in large part due to low cell numbers. *Fgf14* was the most downregulated gene in H3.3K4M Hom tanycytes (Figure S8L-M). Notably, FGF signaling in tanycytes regulates their morphology and proliferation^89,92^, lipid homeostasis^93^ and numerous hypothalamic metabolic process^94,95^.

In sum, perturbation of H3K4 methylation disrupts the organization of tanycytes, astrocytes and oligodendrocyte in the hypothalamus, and specifically the ME. Disruption of this cellular architecture at this critical blood-barrier junction will alter hypothalamic function and hormonal regulation that is likely driving initial growth retardation followed by obesity, increased leptin levels, and contributing to altered behavioral phenotypes in H3.3K4M Hom mice.

## DISCUSSION

Dysregulation of H3K4 methylation is associated with ASD, schizophrenia and numerous other NDDs with ID^8^. While several studies have utilized lysine-to-methionine (K-to-M) mutations to study histone modifications^96^, the Cre-dependent H3 variant H3.3K4M transgenic mouse used here^38^ permits precise spatial and temporal control to study H3K4 methylation during neurodevelopment. We used *Nkx2.1-Cre* mice to express this mutant histone in the embryonic MGE and hypothalamus, two brain regions associated with many H3K4 disease-related phenotypes such as ID, epilepsy and abnormal body growth^3,4^. H3.3K4M Hom mice contain both WT and mutant H3.3 alleles, mimicking the hypomorphs often found in human disease. While ∼50% of H3.3K4M Hom die by 20 weeks, the surviving mice displayed a complex array of deficits observed in many NDDs.

H3.3K4M Hom mice had fewer MGE-derived interneurons in the cortex and hippocampus, particularly PV+ interneurons, which is predicted by the decrease of PV-fated genes *Maf* and *Mef2c* in the MGE. This decrease in interneurons, in combination with altered electrophysiological properties reminiscent of immature interneurons, results in increased epileptiform activity and seizure susceptibility. Our single cell transcriptome analysis identified genes regulating potassium ion flow were significantly downregulated in PV+ cortical interneurons in the H3.3K4M Hom mouse. Notably, perturbation of potassium ion channels and transporters are well-established causes of epilepsy^97,98^, and potassium channels are critical regulators of gamma oscillations^99,100^. Many of these genes were also enriched in the top downregulation co-expression module of PV population in H3.3K4M Hom mice. This finding, in combination with proportional changes of specific PV+ subtypes, indicates that distinct cohorts of genes in certain PV+ interneuron subtypes are preferentially affected upon perturbation of H3K4 methylation.

Mutations that alter histone modifications can have differential effects based on cell types, development stage, and resistance or susceptibility at specific genomic loci^47,96,101^. Consistent with these observations, we found differential effects of perturbed H3K4 methylation on the MGE and hypothalamus. The overwhelming majority of DEGs were downregulated in the MGE of H3.3K4M Hom mice (as were differentially accessible peaks), and this trend was maintained in PV+ and SST+ cortical interneurons. DEGs in the embryonic hypothalamus were more evenly distributed between upregulated and downregulated in the H3.3K4M Hom mouse, while there was a strong bias for differentially accessible peaks being enriched in the mutant hypothalamus. Despite these different trends of upregulated and downregulated DEGs in H3.3K4M Hom mice, the resulting neurogenic phenotype is similar because we observed a decrease in transcription factors driving postmitotic differentiation in the MGE and an increase in genes promoting proliferation in the hypothalamus. This decrease in neurogenic genes and increase in NPC proliferative genes was also observed in the cortex *H3f3a/H3f3b* double KO mice^12^.

What mechanisms could underlie these differential effects between the MGE and hypothalamus? First, K-to-M mutations strongly suppress but do not fully eliminate histone methylation^96^. There is evidence that critical fate-determining genes may be resistant to K-to-M mutations^102^ and other genetic manipulations that target histone modifications^47^. This innate resistance of certain genomic loci likely varies between cell types. Second, H3K4me3 can alter chromatin state in trans by recruiting ATP-dependent chromatin-remodeling complexes, leading to opening of local chromatin and allowing gene transcription at distinct loci in different cells^103,104^. Third, the H3K4M mutation also has a strong binding affinity to several demethylases that target H3K4^105^, which could differentially exacerbate this loss of methylation in distinct cell types. Fourth, in addition to blocking H3K4 methylation, the H3K4M mutation can lead to decreased acetylation of H3K27 (H3K27ac) and decreased protein levels of Mll3/KMT2C and Mll4/KMT2C, two methyltransferases that target H3K4^14,38^. Haploinsufficiency of *MLL4* in humans results in Kabuki syndrome, whose symptoms include short stature, ID and microcephaly^106,107^. In *Mll4*^+/-^ mice, growth hormone-releasing hormone (GHRH)-neurons in ARC are particularly affected because ∼90% of these neurons express *Mll4*, whereas a much lower proportion of other ARC and other hypothalamic neurons express *Mll4*^108^. Thus, depletion of Mll3 and/or Mll4 in H3.3K4M Hom mice would affect growth-regulating cells in the hypothalamus more so than other cell types.

ARC integrates hormonal and nutrient signals to regulate feeding and metabolism^109^. It receives information through from the ME, a circumventricular organ, as well as direct input from tanycytes lining the third ventricle^110,111^. Leptin-sensing ARC neurons project to the PVN to promote or inhibit food intake^112^. Serum leptin levels are higher in the satiated state and are correlated with body fat weight^113^, while an increase in leptin levels along with obesity in the H3.3K4M Hom mice indicates leptin resistance in these mice, which found in most obese individuals^114^. Notably, we observed the largest downregulation of ARC- and PVN-fated neurons in embryonic hypothalamus of H3.3K4M Hom mice compared to other hypothalamic nuclei. Although the RNA level of neuronal ARC markers *Pomc*, *Agrp* and *Npy* were not changed in the P60 hypothalamus of H3.3K4M mice, further work is needed to explore distinct cellular changes in specific hypothalamic nuclei.

Hypothalamic astrocytes also play important roles in energy metabolism by sensing and transporting nutrients in the interface blood vessels and neurons^44^. We observed an increase of hypothalamic astrocytes in H3.3K4M Hom mice, as well as a clear segregation of WT and Hom astrocytes that was not observed in any other cell type. Astrocyte DEGs are associated with cell adhesion, lipid metabolic processes, and glutamate or GABA receptor function, suggesting disruption of multiple astrocytic functions in mutant mice. Additionally, we observed a striking disorganization and altered morphology of tanycytes and glia cells in the hypothalamus of H3.3K4M mice. Tanycytes can act as neural progenitor cells generating new neurons and astrocytes^87^, and they control leptin transport into the third ventricle^115^. The greatest tanycytes DEG is *Fgf14*, and notably, FGF signaling in tanycytes is involved in numerous hypothalamic metabolic process^87,93–95^. Taken together, these cellular and genetic changes likely underlie the increased food intake and fat weight gain leading to obesity observed in the H3.3K4M Hom mice. A similar growth trajectory of initially underweight mice eventually developing obesity was observed in another H3 point mutation (H3.3G34W)^101^. This could indicate a particular sensitivity of the hypothalamic feeding circuit to manipulation of histone modifications, which may explain why so many diseases related to altered histone modifications demonstrate short stature and other growth defects^3,4^.

Mouse models using different strategies to genetically perturb H3K4 methylation are associated with abnormal behaviors, including deficits in learning and memory, impaired sociability, severe hyperactivity, ASD-like behaviors, ID and epilepsy^9,21–23,116^. Here, the H3.3K4M Hom mice display abnormal locomotor and gait activity, increased anxiety, impaired spatial memory and sensorimotor gating, hypoactivity and impulsive activity (ADHD-like phenotypes) compared with WT. It is well-established that many NDDs affect one sex more than the other^104,117–120^. We found that H3.3K4M Hom female mice displayed more severe deficits in many of these behaviors compared to males, likely due in part to a decrease of potassium ion channels in Hom female compared to Hom male of PV population. While males and females have a similar prevalence of having epilepsy, females have greater risk for generalized-onset epilepsies, such as juvenile myoclonic epilepsy and juvenile absence epilepsy, with differential sex hormone control by the hypothalamus as one strong candidate driving these differences^121^. Additionally, there is evidence for differential developmental expression and function GABA-related signaling between males and females that may contribute to increased female susceptibility in H3.3K4M Hom mice^122,123^.

NDDs such as autism, schizophrenia, ADHD, ID, and various learning and motor disorders are more prominently diagnosed in males^117,124^, whereas obsessive-compulsive-like tic behaviors, anxiety, bipolar disorder, depression and personality disorders are more prominent in females^125,126^. Female H3.3K4M Hom mice displayed greater deficits in numerous motor-related assays compared to males. Stimulation of Nkx2.1+ neurons in the ventromedial hypothalamus (VMH) generated a female-specific increase in movement, and loss of these cells resulted in females becoming obese, but not males^127^. Additionally, the MGE gives rise to interneurons and GABAergic projection neurons that populate the basal ganglia, a structure critical for voluntary movements which shows sexually dimorphism in humans^128,129^, and specifically with MGE-derived interneuron distribution in the striatum^130^. The preoptic area in the hypothalamus and the bed nucleus of the stria terminalis (BNST), both of which contain Nkx2.1-lineage cells, display sexual dimorphism^131^. Notably, these regions also display sex differences in H3K4me3 levels, with ∼70% of differential peaks enriched in females, and many at genomic loci associated with seizures, emotion/affective behavior, and learning and memory^120^. Additionally, H3K4me is correlated with escape from X inactivation^132^. Thus, a bias for H3.3K4M Hom female-enriched deficits in various locomotor and behavior assays likely arises from differential transcriptome, cellular composition and circuit alterations in the hypothalamus and MGE-derived GABAergic cells.

In sum, reduction of H3K4 methylation in the embryonic MGE and hypothalamus recapitulates the most consistent phenotypes observed in H3K4-associated diseases: epilepsy, altered cognitive function, and abnormal body growth. H3.3K4M Hom mice had fewer interneurons, reduced network synchrony, and downregulation of many genes in PV+ interneurons associated with potassium ion transport, ultimately leading to increased seizure activity in these mice. In the hypothalamus, there was a significant change in the relative proportion of cells and striking cellular disorganization along the blood-brain barrier. These changes, combined with altered transcriptome profiles, likely disrupts normal hormonal regulation, energy metabolism and other critical hypothalamic functions leading to altered body growth and behavioral phenotypes in these mice.

## METHODS

### Animals

All experimental procedures were conducted in accordance with the National Institutes of Health guidelines and were approved by the NICHD Animal Care and Use Committee (protocol #23-047). The following mouse lines were used in this study: *LSL-K4M* (gift from Dr. Kai Ge, NIDDK)^38^; *Nkx2.1-Cre* (Jax# 008661)^41^, *Ai9* (Jax# 007909)^133^, *Sun1-sfGFP* (Jax# 030952)^134^. *LSL-K4M* mice were genotyped with forward primer (5’-GGGTCTGTTCGAAGATACCAACC-3’) and reverse primer (5’-GTAGTCGGGCACGTCGTA-3’), with heterozygous and homozygous mice determined via qPCR. Mice were housed under standard conditions (12h light and 12h dark). The morning on which the vaginal plug was observed was denoted E0.5.

### Brain harvesting

Embryonic brain dissections: Pregnant dams were anesthetized with an i.p. injection of Euthasol (270 mg/kg, 50 µL injection per 30 g mouse), E13.5 embryos were removed and placed in ice-cold carbogenated artificial cerebral spinal fluid (ACSF, in mM: 87 NaCl, 26 NaHCO_3_, 2.5 KCl, 1.25 NaH_2_PO_4_, 0.5 CaCl_2_, 7 MgCl_2_, 10 glucose, 75 sucrose, saturated with 95% O_2_, 5% CO_2_, pH 7.4). Embryonic tails were cut for genotyping. MGE and hypothalamus were dissected from individual brains in ice-cold ACSF, transferred to individual 0.5 mL Eppendorf tubes, then flash frozen on dry ice and stored at -80°C. For analyzing fixed tissue, E13.5 brains were removed and drop-fixed in 4% paraformaldehyde (PFA) in PBS overnight, washed in PBS, incubated in 30% sucrose in PBS at 4°C overnight, embedded in OCT and stored at -80°C.

Adult brain dissections: P21-P150 mice were anesthetized with Euthasol. For harvesting brain regions, the somatosensory cortex and hypothalamus were dissected, transferred to 1.5 mL Eppendorf tubes, then flash frozen on dry ice and stored at -80°C. For analyzing fixed brains, mice were perfused with PBS followed by 4% PFA, brains were removed and post-fixed in 4% PFA overnight, washed in PBS, incubated in 30% sucrose in PBS at 4°C overnight, then embedded in OCT and stored at -80°C.

### Tissue staining and microscopy

P21-P150 brains were cryosectioned at 30 µm, sections transferred to 96-well plates containing antifreeze solution (30% ethylene glycol, 30% glycerol, 40% PBS) and stored at -20°C. Prior to staining, brain sections were washed in PBS (3 x 10 minutes). Brain sections were incubated in block solution (10% Normal Donkey Serum in PBS + 0.3% Triton X-100) for 1 hour RT, then incubated for 24-48 hour in primary antibodies in block solution at 4°C. Then sections were washed in PBS (4 x 10 minutes) and incubated with secondary antibodies and DAPI for 2 hours at RT or overnight at 4°C. Brain sections were washed and mounted.

E13.5 embryonic brains were cryosectioned at 14 µm, mounted on slides, dried and stored at -20°C. Slides were incubated in block solution for 1 hour RT, then incubated overnight with primary antibodies in block solution and secondary antibodies for 1-2 hours RT.

The following primary antibodies were used in this study: rat-anti SST (1:300, Millipore MAB354), goat-anti PV (1:1000, Swant PVG213), rabbit-anti nNos (1:500, Millipore MAB5380), rabbit-anti H3K4me3 (1:500, Cell Signaling TECHNOLOGY 9751S), rat-anti HA (1:200, Roche 11867423001), mouse-anti GFAP (1:500, Sigma-Aldrich G3893), rabbit-anti NeuN (1:500, abcam ab177487), rabbit-anti Vimentin (1:500, Cell Signaling TECHNOLOGY 5741T), rabbit-anti Olig2 (1:500, Sigma-Aldrich AB9610), mouse-anti MBP (1:1000, Invitrogen MA1-10837). Species-specific fluorescent secondary antibodies (1:500) were used conjugated to AlexaFluor 488, 647 and 790.

Nissl Staining: Slides were warmed to RT and then treated in the following solution sequence for 5 minutes each: 70% EtOH, 95% EtOH, 100% EtOH, 95% EtOH, 70% EtOH and 50% EtOH. Slides were then rinsed with ddH_2_O, incubated in 0.1% warm crystal violet stain for 10-20 min, then washed with ddH_2_O. Slides were then incubated in 70% EtOH and 95% EtOH for 5 minutes each, then 100% EtOH for 2 minutes twice. Slides were then incubated with a xylene substitute for 5 minutes twice, air dried overnight and mounted with coverslips.

Fluorescent in situ Hybridization: Brains were cryosectioned at 14 µm. RNAscope HiPlex and HiPlexUp in situ hybridization assays (Advanced Cell Diagnostics) were performed according to the manufacturer’s instructions. All probes and reagents from Advanced Cell Diagnostics: probe-Mm-Reln-C3 (405981-C3), probe-Mm-Pvalb (421931) and tdtomato-C2 (317041-C2).

All images were captured on an Olympus VS200 scanner (VS200 ASW) or Zeiss Axioimager.M2 (with Zen Blue software). Post-processing was performed with Adobe Photoshop or ImageJ.

### Electrophysiology experiments

Adult mice were anesthetized with isoflurane and decapitated. Brains were quickly dissected and placed in ice-cold artificial cerebrospinal fluid (ACSF) modified for dissection containing (in mM): 80 NaCl, 25 NaHCO_3_, 10 glucose, 1.25 NaH_2_PO_4_, 3.5 KCl, 4.5 MgCl2, 0.5 CaCl_2_, 90 sucrose (Sigma-Aldrich) and bubbled with carbogen (95% O_2_ and 5% CO_2_). Horizontal hippocampal sections were prepared from the two hemispheres using a Leica VT1200S vibratome (Leica Microsystems). Recordings were performed using a Multiclamp 700B amplifier and acquired using pCLAMP 10,4 software (Molecular Devices). Signals were acquired at 5 kHz, digitized, and stored using Digidata 1322A and pCLAMP 10,4 software (Molecular Devices, CA, USA).

### Ex-vivo gamma oscillation recordings and analysis

Nine 3-month-old mice were used for local field potential (LFP) recordings (WT n=4 and Hom n=5 with both male and female). 350 µm thick hippocampal slices were placed into an interface holding chamber containing ACSF modified for LFP: 124 mM NaCl, 30 mM NaHCO_3_, 10 mM glucose, 1.25 mM NaH_2_PO_4_, 3.5 mM KCl, 1.5 mM MgCl_2_, 1.5 mM CaCl_2_ (Sigma-Aldrich) continuously supplied with humidified carbogen gas (95% O_2_ and 5% CO_2_). The chamber was held at 34°C during the slicing process and subsequently allowed to cool down to room temperature (∼22°C) for at least 1 h in order to let the slices recover. Hippocampal gamma oscillations experiments were performed as described previously^135^. Oscillations were recorded at 34°C using borosilicate microelectrodes (1.5–2.5 MΩ) filled with ACSF placed in the CA3 stratum pyramidale and elicited by adding kainic acid (400 nM; Tocris Bioscience) to the extracellular bath. The oscillations were allowed to stabilize for at least 20 min before recordings were performed. Signals were conditioned using a HumBug 50 Hz noise eliminator (Quest Scientific).

Before analysis, signals were low pass filtered at 0.1 KHz and high pass filtered at 10 Hz. For oscillation power spectra, Fast Fourier Transformations were obtained from 60s of LFP recording (segment length 8192 points). Frequency variance data was obtained from the power spectra described above using the Axograph software (Kagi, Berkeley, CA, USA). Gamma power was calculated by integrating the power spectral density from 20 to 80 Hz using Clampfit 11.2.

### Whole-cell patch clamp recordings and analysis

Six 8-week-old mice (n=3 WT and Hom female brains) were used to perform whole cell patch clamp recordings of MGE derived hippocampal interneurons. 300 µm thick hippocampal slices were immediately placed into a submerged holding chamber containing standard ACSF: 130 mM NaCl, 24 mM NaHCO_3_, 10 mM glucose, 1.25 mM NaH_2_PO_4_, 3.5 mM KCl, 1.5 mM MgCl_2_, 2.5 mM CaCl_2_ (Sigma-Aldrich) continuously supplied with humidified carbogen gas (95% O_2_ and 5% CO_2_). Whole-cell patch clamp recordings were performed on slices in the submerged recording chamber at 34°C with borosilicate glass electrodes filled with intracellular recording solution containing 130 mM K-gluconate, 5 mM KCl, 10 mM HEPES, 3 mM MgCl_2_,2 mM Na_2_ATP, 0.3 mM NaGTP, 0.6 mM EGTA, and 0.2% biocytin. Voltage clamp and current clamp configuration were used.

Analysis of whole cell patch clamp recordings was performed using the AutoAnt Software^136^. Principal Component Analysis (PCA) was used to classify 93 interneurons (WT n=49 and Hom n=44) based in their intrinsic properties (Input resistance, membrane time constant, membrane capacitance, sag ratio, maximum firing rate and action potential half-width). The PCA was performed in Python with the Sklearn package. The kmeans clustering algorithm was used to divide the standardized data using 3 principal components. The adequate number of clusters for the dataset was identified using the Silhouette Score.

### Pentylenetetrazole (PTZ) injection and recording

PTZ was prepared on the day of use with in sterile 0.9% (w/v) NaCl. PTZ injection and recording was performed on 3-month and 5-month-old mice. All 5-month animals will receive an IP injection of PTZ (Sigma-Aldrich, P6500) at 20 mg/kg and were recorded immediately via video tracking system ANY-maze (Stoelting Company) for 60 min. 3-month female animals will receive an IP injection of PTZ at 20 mg/kg, while 3-month male animals will receive an IP injection of PTZ at 40 mg/kg due to less sensitive to PTZ. Classify and score the epileptic behaviors based on previous protocol^137^ using the following criteria: stage 0 = normal behavior, stage 1 = immobility and rigidity; stage 2 = head bobbing, facial, forelimb, or hindlimb myoclonus; stage 3 = continuous whole-body myoclonus, myoclonic jerks or tail held up stiffly; stage 4 = tonic seizure, continuous rearing and falling, stage 5 = clonic-tonic seizure, stage 6 = death. Total seizure scores were calculated at every 5-min block in the first 20 min. The mortality was also noted during the recording process.

### Behavioral tests

All behavioral tests were performed on WT, Het and Hom littermates according to approved NICHD and/or NIMH standard operating procedures (SOPs) and cleaning protocols. Animal behavioral tests were carried out during the daylight cycle in NICHD animal behavioral room or at the NIMH Rodent Behavior Core. All mice were habituated to the testing environment for 30-60 minutes before behavior tests. All behavior assays were performed blind to genotype and 1-3 independent cohorts were used for each behavioral test. The sequence of tests was as follows, with at least one week elapsed between any two behavior tests. Juvenile and young adults (6-11 weeks old): Home-cage test (5-6 weeks old); Open-field test (7-8 weeks old); Three-chamber test (9-10 weeks old); Barnes maze test (10-11 weeks). Adults (3-5 months old): Free-walk test; Light-dark box test; Novel object recognition test; Cliff avoidance reaction; Prepulse inhibition startle test.

### Home-cage test

Individually housed juvenile mice were observed in their home cages placed in the Photobeam Activity System-Home Cage (PAS-HC, SD Instruments) to assess activity (ambulation, fine movements and rearing) that can be quantified by beam breaks. Mice were kept in the cage for 4 days with appropriate food and water. During this testing, mice were housed in a ‘quiet’ room to minimize outside interactions. Mice were returned to group housing after experiment. The total ambulatory movements, fine movements, rearing and general locomotor activity were analyzed and processed using the PAS-HC data collection transferred to Excel. N (female) = 8 WT and 11 Hom; N (male) = 7 WT and 4 Hom.

### Open-field test

Mice were originally placed in the center of open field apparatus (40 cm x 40 cm x 30 cm Plexiglas box) and recorded for 25 minutes. The average speed and time spent in the different open field areas were recorded and analyzed with the video tracking system ANY-maze (Stoelting Company). N (female) = 13 WT, 8 Het and 11 Hom; N (male) = 12 WT, 11 Het and 7 Hom.

### Three-chamber test

Tests were performed in an opaque Plexiglas rectangular box (60 cm x 45 cm x 20 cm) consisting of 3-chambers, with clear dividing walls and an open central chamber allowing free access between chambers. Wire cups were used to hold a single stimuli mouse (C57BL/6J mice that were sex and age-matched with test mice). The test consists of three 10-minute phases that were all recorded using the ANY-maze system. For phase 1, the test mouse was placed in the center chamber with closed walls. For phase 2, two empty wire cups were placed in each side chambers and the closed walls were opened, which let the mouse explore all three chambers. For phase 3, the test mouse was confined in the center chamber and a stimuli mouse was placed in a wire cup on one side of the chamber. Both walls were removed and the test mouse explored chambers for 10 minutes. Sniffing (investigating) was defined as being within 3 cm of either cup’s edge. The total time spent in each region and time investigating each stimulus (stimuli mouse or empty cup) was recorded and quantified. N (female) = 13 WT, 6 Het and 11 Hom; N (male) = 12 WT, 11 Het and 7 Hom.

### Barnes maze test

The Barnes maze consists of a white elevated circular platform (diameter = 122 cm) with 80 cm support frame containing 20 equally distributed holes (diameter = 5 cm), with 1 ‘escape’ hole leading to a hidden black drop box. Light source was affixed above the platform to brightly illuminate the maze, and 90 dB white noise generated from loudspeakers was used to motivate mice to enter the dark escape box. Three distal visual cues surrounded the platform.

The protocol used a 6-day paradigm that included a habituation, training and probe trial phase. During habituation (day 1), the mouse was placed in the escape tunnel for 1 min and then placed in the center of platform until it enters the escape hole or 3 minutes elapse. In the training days (days 2-5), the mouse is placed in a start chamber located in the center of one of the four quadrants and the mouse explores the platform until it enters the escape hole or 3 minutes elapse. If a mouse fails to find the escape tunnel within 3 minutes, the mouse is guided to the dark escape tunnel and remains there for 15 seconds. Mice underwent 4 trials on each training day. For the probe trial (day 6), the escape tunnel is removed and mice are placed in the center of the platform to explore the maze for 90 seconds. Time spent around target hole was defined as being within 1 cm of target hole edge. All trials were recorded using the ANY-maze system allowing quantification of time each mouse spent in the target hole and target quadrant. N (female) = 13 WT and 9 Hom; N (male) = 8 WT and 4 Hom.

### Free-walk test

FreeWalkScan 2.0 (CleverSys Inc.) was used to assess mice gait. 3-month-old mice with similar body weight move freely in a 40 cm × 40 cm × 30 cm (length × width × height) chamber. A high-speed camera below a clear bottom plate captures mouse movement for 5 minutes with red light in dark room. Videos are analyzed using FreewalkScanTM 2.0 software for various characteristic of gait, including BaseFront (distance between sequential footprints of front limbs), StrideLength (distance between two sequential footprints of same paw), Contact Size, Average Pressure and Stance Time (time corresponding paw stays on ground) of each paw. N (female) = 9 WT, 9 Het and 4 Hom; N (male) = 6 WT, 9 Het and 8 Hom.

### Light-dark box test

The apparatus (40 cm x 40 cm x 30 cm) consists of dark chamber (40 cm x 15 cm x 15 cm) and light chamber compartment. There is a door between two chambers which allow mice to freely move between chambers. Bright white lights illuminated the box from above as mice were placed in the box for 10 minutes. The test was recorded by ANY-maze video trace camera to determine the amount of time each mouse spent in the light chamber. N (female) = 14 WT, 12 Het and 7 Hom; N (male) = 9 WT, 11 Het and 9 Hom.

### 2 Object Novel object recognition test

Novel object recognition test was conducted in an open field arena (40 cm x 40 cm white Plexiglas box). The protocol used a 3-day paradigm that includes habituation, training, and testing phase. During habituation (day 1), mice were placed into the center of an open field arena and allowed to explore for 10 minutes. For training (day 2), two identical objects were placed on either side of the center of the arena, and the mice explored the arena for 10 minutes. For testing (day 3), a familiar object and a novel object (dissimilar object) were placed in the same position as in the training day and mice explored the arena for 10 minutes. ANY-maze was used to record each trial. The amount of time each mouse spent with their nose oriented toward the object within 2.5 cm of the object edge was considered ‘exploration time’.

The recognition index was calculated as: Recognition index = (time spent exploring novel − time spent exploring familiar)/(total time spent exploring both objects). N (female) = 9 WT, 10 Het and 7 Hom; N (male) = 6 WT, 7 Het and 6 Hom.

### Prepulse inhibition (PPI) of the acoustic startle reflex

Startle response and pre-pulse inhibition were performed via SR-LAB-Startle Response System (San Diego Instruments). Mice are placed in a startle apparatus with sound-attenuated chamber. Whole-body startle movements were detected by a piezoelectric accelerometer mounted beneath the cage, converted to electrical signals, and digitized and stored by a computer. A loudspeaker mounted to the side of the cage produced white noise acoustic stimuli. A continuous 64 dB background noise was present throughout the test. The test began with a 5-minute acclimation to the apparatus, followed by 12 pulse-alone trials. To measure PPI, 12 blocks containing 6 trials were presented: 1 pulse-alone trial, 4 prepulse-pulse trials, and 1 no-stimulus trial. Pulse-alone trials consisted of a 40 ms startle pulse at 120 dB; prepulse-pulse trials consisted of a 20 ms prepulse at 66, 68, 72, or 76 dB followed 100 ms later by the pulse; and no-stimulus trials consisted of background noise only. Trials will be presented in pseudorandom order, with an inter-trial interval of 10-20 s (mean of 15 s). Startle response will be determined as the peak amplitude within 100 ms of the startle pulse onset. The entire test lasted 30 minutes. N (female) = 10 WT and 7 Hom; N (male) = 6 WT and 5 Hom.

### Cliff Avoidance Reaction (CAR) Test

The CAR test was performed as protocol^65^ with litter modified to assess impulsive-like behaviors. The apparatus consists of a square platform (20 cm x 20 cm) and support by a rod (Height = 40cm) that was placed in an open field box with soft padding. After habitation, the mouse was gently placed onto the platform, and they were also placed back onto platform after falling. The latency to first jump off and numbers of jump off were recorded. The test was performed and analyzed for 15 minutes under white lights. N (female) = 13 WT and 7 Hom; N (male) = 5 WT and 5 Hom.

### Body composition and Elisa analysis

Analysis of body composition in live mice using EchoMRI100 analyzer (Echo Medical Systems) in the Mouse Metabolism Core of NIDDK. This test revealed both the fat weight and lean weight for each mouse. Serum leptin levels were measured using a mouse leptin ELISA kit (ThermoFisher, #KMC2281) in accordance with their standard protocol.

### Cell counting and analysis

All cell counts on *Nkx2.1-Cre*;*K4M*;*Ai9* mice were performed manually using Photoshop and were blind to genotypes. Cortex: Cells were counted from 3 non-consecutive brain sections through the somatosensory cortex for each P21 brain, from 5 WT, 4 Het and 5 Hom mice. DAPI staining was used to divide the cortex into superficial (I-III) and deep (IV-VI) layers. Tom+, PV+ and SST+ cells were counted and the density was calculated by cells/mm^2^. Any PV+ and SST+ cells that were Tom-were excluded from the counts. Hippocampus: Cells were counted from 4 non-consecutive sections spanning the anterior-to-middle P21 dorsal hippocampus, from 5 WT and 5 Hom brains. Tom+, PV+, SST+ and nNOS+ cells counted and the density was calculated by cell numbers/mm^2^. Small Tom+ cell bodies in CA2/3 that are primarily Olig2+ oligodendrocytes were excluded from these interneuron counts and counted as separate group. Hypothalamus: Six coronal brain sections (bregma: from -1.58 to -1.94 mm) were analyzed per mouse. The mean fluorescent intensity of GFAP in region of interest was measured using ImageJ software (v1.53; NIH) via from 4 WT and 3 Hom of adult mice (3-5 months old).

### Western blot

Total core histone proteins were extracted using the EpiQuik Total Histone Extraction Kit (EpigenTek). E13.5 MGE and hypothalamus samples were obtained as described above. Protein amount was qualitied using Pierce BCA Protein Assay Kit (ThermoScientific), typically ∼35 µg protein was obtained from tissue from 1 embryo. Total 40 µL Loading samples were prepared with Sample Reducing Agent (ThermoFisher, B0009) and Bolt LDS Sample Buffer (ThermoFisher, B0007), and were load on a 4-12% Bolt Bis-Tris Plus Mini Protein Gel (ThermoFisher, NW04120BOX), running 20-30 minutes at 200V with Blot MES SDS Running buffer (ThermoFisher, B0002). After that, gel was transferred to PVDF membranes using the iBlot 2 Gel Transfer Device (Invitrogen) and transferred for 7 min at 20V. Membrane was blocked 30 min in block buffer (SuperBlock Dry Blend Blocking Buffer, ThermoFisher #37545). Membranes were incubated in primary antibody overnight at 4°C, secondary antibody for 1 hour at RT, washed 3 times with TBST washing buffer, and then blots were imaged on the ChemiDoC MP Imaging System (Bio-Rad). The following primary antibodies were used: mouse anti-H3 (1:1000, Cell Signaling TECHNOLOGY #3638), rabbit anti-H3K4me3 (1:500, Cell Signaling Technology #9751), rabbit anti-H3K4me1 (1:1000, Abcam ab8895), rabbit anti-H3K4me2 (1:500, Abcam ab7766). The following secondary antibodies were used: anti-mouse-Starbright Blue-520 (Bio-Rad# 64456855; 1:2000) and anti-Rabbit-Starbright Blue-700 (Bio-Rad# 64484700; 1:2000).

### Multiome sequencing and analysis

Frozen E13.5 and P60 tissue from *Nkx2.1-Cre*;*H3.3K4M;sun1-GFP* mice were mechanically lysed, washed and filtered to generate single nuclei suspensions for the 10x Genomics Multiome assay as previously described^138^. These suspensions were then sorted via flow cytometry (Sony SH800) to harvest ∼100,000 GFP+ nuclei. Nuclei solutions were centrifuged in a bucket centrifuge at 500 g for 5 minutes at 4°C. Supernatant was removed and nuclei reconstituted in 1x Nuclei buffer at a density of 3,000 nuclei/µl. 5 µl of this solution was then used for the 10x Genomics Single Cell Multiome Assay per manufacturer’s protocol. Our single cell Multiome workflow can be found at https://nichd-bspc.github.io/multiome-wf/. Samples details and sequencing information are described in Supplementary Table 1.

Sequencing analysis: Gene expression and open chromatin state were profiled using 10X Chromium Single Cell Multiome ATAC + Gene Expression kit (10X Genomics, 1000285). Joint libraries were simultaneously created by following standard protocol. Sequencing was conducted with paired-end (50 x 50 bp) using an Illumina HiSeq 2500 or NovaSeq 6000. The raw sequencing data were processed using Cell Ranger ARC (v2.0.0) pipeline. The cellranger-arc mkfastq command was used to generate the demultiplexed FASTQ files from BCL files. The sequencing reads were aligned to custom built mouse (GRCm38/mm10) reference genome using cellranger-arc count. The cellranger-arc count command was used to generate gene-by-barcode (snRNA-seq) and peak-by-barcode (snATAC-seq) matrices. The cellranger-arc aggr command, without depth normalization (--normalize = none) was used to aggregate sample datasets into a single feature-barcode matrix file.

snRNA-seq data analysis: The aggregated feature-barcode matrix was used as input to Seurat (v5.2.1)^139^ in R (v4.4.2, https://cran.r-project.org). Low-quality cells were removed based on the following QC metrics: RNA feature counts, UMI counts, mitochondrial counts and ribosomal counts. Cells that have RNA feature counts more than 100 and UMI counts less than three standard deviations (SD) of the mean were kept. Cells that have less than 5% mitochondrial counts and 10% ribosomal counts were kept. In addition, R package scDblFinder (v1.20.0)^140^ was used to remove the computationally likely doublets. After filtering out low-quality cells and doublets, we performed standard Seurat workflow with default parameters unless otherwise noted: NormalizeData (LogNormalize), FindVariableFeatures (nfeatures = 2000), ScaleData, RunPCA, FindNeighbors, FindClusters and RunUMAP steps. Dim was set to 30 based on inspection of variance across the principal components (PCs) using the ElbowPlot function.

snATAC-seq data analysis: The aggregated peak-by-barcode matrix was used as input to Signac (v1.14.0)^141^ in R. Low-quality cells were removed based on the following QC metrics: number of chromatin accessibility peaks, nucleosome signal and transcription start site enrichment score. Cells with more than 1000 chromatin accessibility peaks, a transcription start site enrichment score greater than 2, and a nucleosome signal less than 4 were kept. Doublets were also removed by using scDblFinder (v1.20.0)^140^. After filtering out the low-quality cells and doublets, we proceeded with normalization, identified highly variable peaks and reduced dimensions using the function of RunTFIDF, FindTopFeatures (min.cutoff = “q0”) and RunSVD in Signac with default parameters. The first Latent Semantic Indexing (LSI) component was not used it downstream analysis due to its strong correlation with the sequencing depth. Then, regular non-linear dimension reduction and clustering were performed by RunUMAP (dims = 2:30), FindNeighbors (dims = 2:30) and FindClusters in Seurat. The gene activity scores were calculated using the GeneActivity function and added to Seurat object.

Multimodal analysis: The shared cells across the two modalities (snRNA-seq and snATACseq datasets) were merged, and the RNA and ATAC dimensionality reduction was recomputed using default parameters. Multimodal data integration was performed by finding the Weighted Nearest Neighbor (WNN)^142^ using Seurat::FindMultiModalNeighbors. The dimensionality reduction was separately set to 1:30 (snRNA assay) and 2:30 (snATAC assay). The joint UMAP and clustering was performed using WNN graph with default parameters.

Subcluster and annotation analysis: To better analyze the different tissue, cells belonging to E13.5 MGE, E13.5 hypothalamus, P60 cortex and P60 hypothalamus from total Multiome data were separately subsetted. Shared cells across the two modalities were merged, and the RNA and ATAC dimensionality reduction was recomputed using default parameters. Multimodal data integration was recomputed by finding the Weighted Nearest Neighbor (WNN)^142^ using Seurat::FindMultiModalNeighbors. Cell types of E13.5 MGE and P60 hypothalamus datasets were annotated with mature cell type biomarkers. Cell types of P60 cortex datasets were annotated via mapping a reference scRNA-seq dataset of adult mouse cortex^143^ to our datasets with TransferData function and cell types with low numbers were removed. Putative hypothalamic nuclei of E13.5 hypothalamus datasets were annotated via gene expression studies^76–78^. In E13.5 hypothalamus datasets, we identified some cells corresponding to the adjacent prethalamus and telencephalon area via region-specific markers^76^, and the telencephalon cells were removed before performing other analysis.

Differential analysis: Differentially expressed genes or peaks between different conditions (genotype or gender) were computed using the Wilcoxon rank sum test implemented in the Seurat FindMarkers function with Bonferroni correction (significant determination: log_2_FC > ± 0.2, adjusted P-value < 10^-6^), and for DEGs in tanycytes (significant determination: adjust P-value <0.05) due to low cell numbers. The EnhancedVolcano function from the EnhancedVolcano package (v1.24.0, https://github.com/kevinblighe/EnhancedVolcano) was used to visualize the DEGs between different conditions per population and parameters.

Enrichment analysis: Gene ontology pathway analysis was performed on each group’s DEGs with adjusted p values <0.05 using the enrichGO function from the clusterProfiler package (v4.14.0)^144^. The parameters were set as following: OrgDb = org.Mm.eg.db (v3.20.0), keyType = “SYMBOL”, ont = “BP”, universe=“All detected genes in specific population”. ClusterProfiler::simplify function was also used to reduce the redundancy among enriched terms.

Co-expression network analysis: The weighted gene co-expression network analysis in high dimensional transcriptomics (hdWGCNA) was performed by the R package hdWGCNA (v0.4.03)^73^. We set up Seurat objects for WGCNA using cells from P60 cortex interneurons with SetupForWGCNA function and constructed metacells in each group with MetacellsByGroups function. The expression matrix of the PV+ population was prepared with SetDatExpr function. The co-expression relationships between genes were raised by the power to focus on strong connections. Thus, the soft power threshold was set to 6 by performing a parameter sweep with TestSoftPowers function, and the co-expression in the PV+ population was constructed with ConstructNetwork function. Module Eigengenes and connectivity were computed with default parameters in PV population. Then, the top hub genes were determined with GetHubGenes function. We ran the UMAP algorithm on hdWGCNA topological overlap matrix with RunModuleUMAP function and visualized the UMAP co-expression network with ModuleUMAPPlot function in hdWGCNA. Differential Module Eigengene (DME) analysis was also performed between different genotypes in the PV population using FindDMEs function (test.use = “wilcox”) and visualized with PlotDMEsVolcano function in hdWGCNA with default parameters.

Pseudo-time analysis: Single-cell trajectory analysis was performed using Monocle 3 (v1.3.7)^145^ workflow. MGE Seurat object was converted to a CellDataSet object using as.cell_data_set function. The reduce dimensionality of CellDataSet object was set to WNN.UMAP of MGE Seurat object. Learning trajectory graph using learn_graph function. The root of trajectory was set to *Nes+* clusters and order cells using order_cells function with default parameters. Plotting trajectory colored by pseudotime with plot_cells function.

Visualization: Uniform Manifold Approximation and Projection (UMAP) coordinates and WNN clustering, computed by Seurat on multimodal integrated datasets, were visualized using DimPlot function. The expression of genes of interest was visualized using FeaturePlot, VlnPlot or DoHeatmap function. Packages ggplot2 (v3.5.1) and patchwork (v1.3.0) were used to visualize during Multiome datasets analysis process.

## ACKNOWLEDGEMENTS

We thank all members of the Unit on Cellular and Molecular Neurodevelopment, as well as Pedro Rocha, Ariel Levine and Kai Ge for discussion and comments on this project and manuscript. We thank Dr. Kai Ge (NIDDK) for the *LSL-K4M* mice. We thank the NICHD Molecular Genomics Core, specifically Fabio Faucz, Vivek Mahadevan, Tianwei Li and James Iben; the NIDDK Mouse Metabolism Core, particularly Oksana Gavrilova and Naili Liu, for assistance with measuring body composition; the NINDS and NICHD animal facility for mouse husbandry assistance. This work utilized the computational resources of the NIH HPC Biowulf cluster (http://hpc.nih.gov). This work was supported, in part, by the NIMH IRP Rodent Behavioral Core (MH002952). This project was funded by NICHD intramural projects HD008962 (T.J.P.), HD008986 (R.K.D.), HDXXXXX (C.J.M.); NICHD Scientific Director’s Award (T.J.P); and an NICHD Intramural Research Fellowship (J.L.). The content of this publication does not necessarily reflect the views or policies of the Department of Health and Human Services, nor does mention of trade names, commercial products or organizations imply endorsement by the U.S. Government.

## DATA AND SOFTWARE AVAILABILITY

All single cell Mulitome sequencing data reported in this paper has been submitted to GEO with the accession number GSE293881 and will be publicly available upon publication. For any additional inquiries about data accessibility and analysis, please.

---

**Figure S1.**
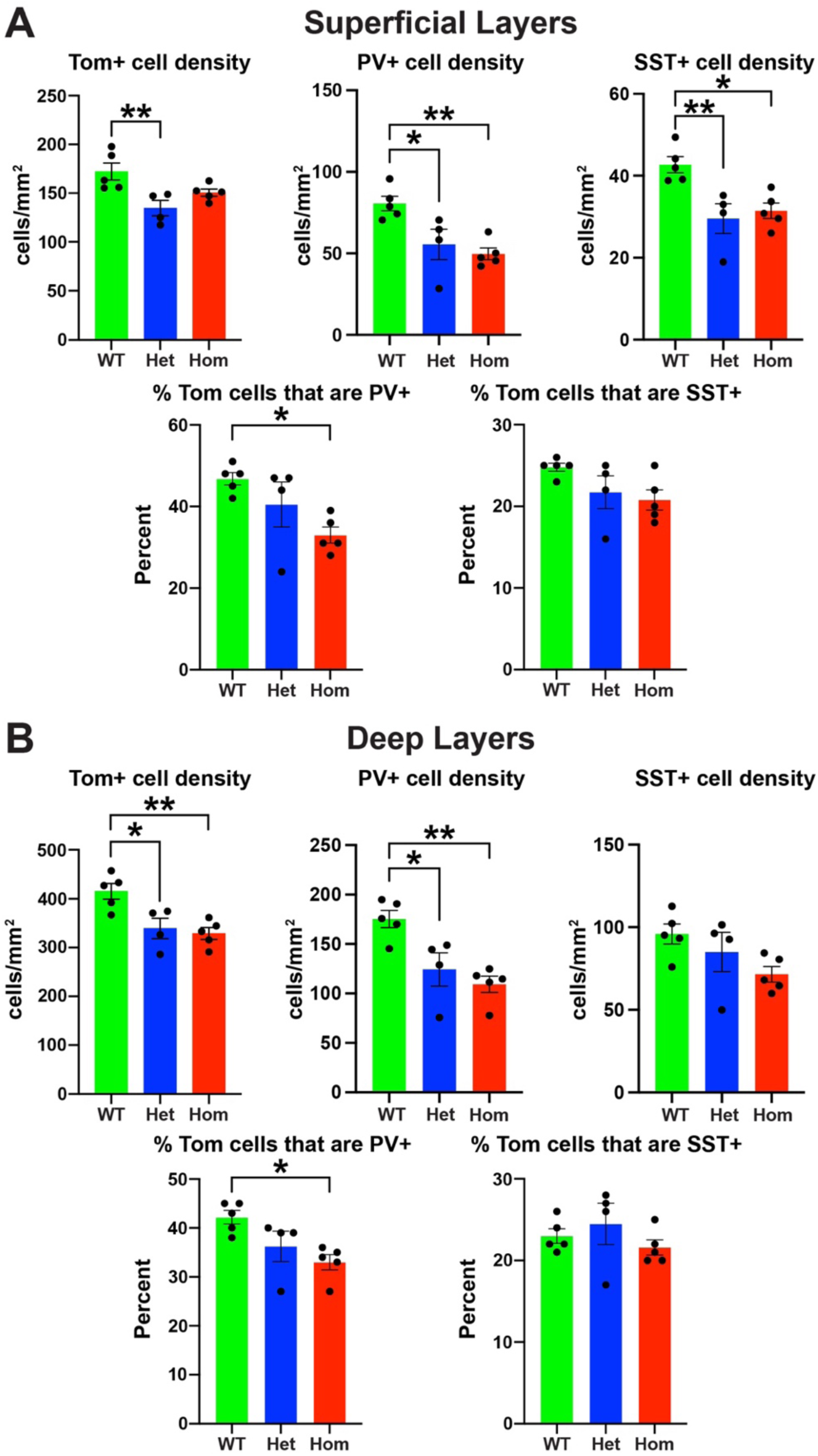
Decreased MGE-derived interneurons in the superficial and deep layers of H3.3K4M Hom mice cortex. **A.** Graphs depicting the density of Tom+, PV+ and SST+ cells (top) and the percent of Tom+ cells expressing PV or SST (bottom) in the superficial layers (I-III) of P21 *Nkx2.1-Cre*;*H3.3K4M*;*Ai9* mice. **B.** Graphs depicting the density of Tom+, PV+ and SST+ cells (top) and the percent of Tom+ cells expressing PV or SST (bottom) in the deep layers (IV-VI) of P21 *Nkx2.1-Cre*;*H3.3K4M*;*Ai9* mice. All stats are one-way ANOVA followed by Tukey’s multiple comparison tests (A, B): * = p ≤ .05, ** = p ≤ .005.

**Figure S2.**
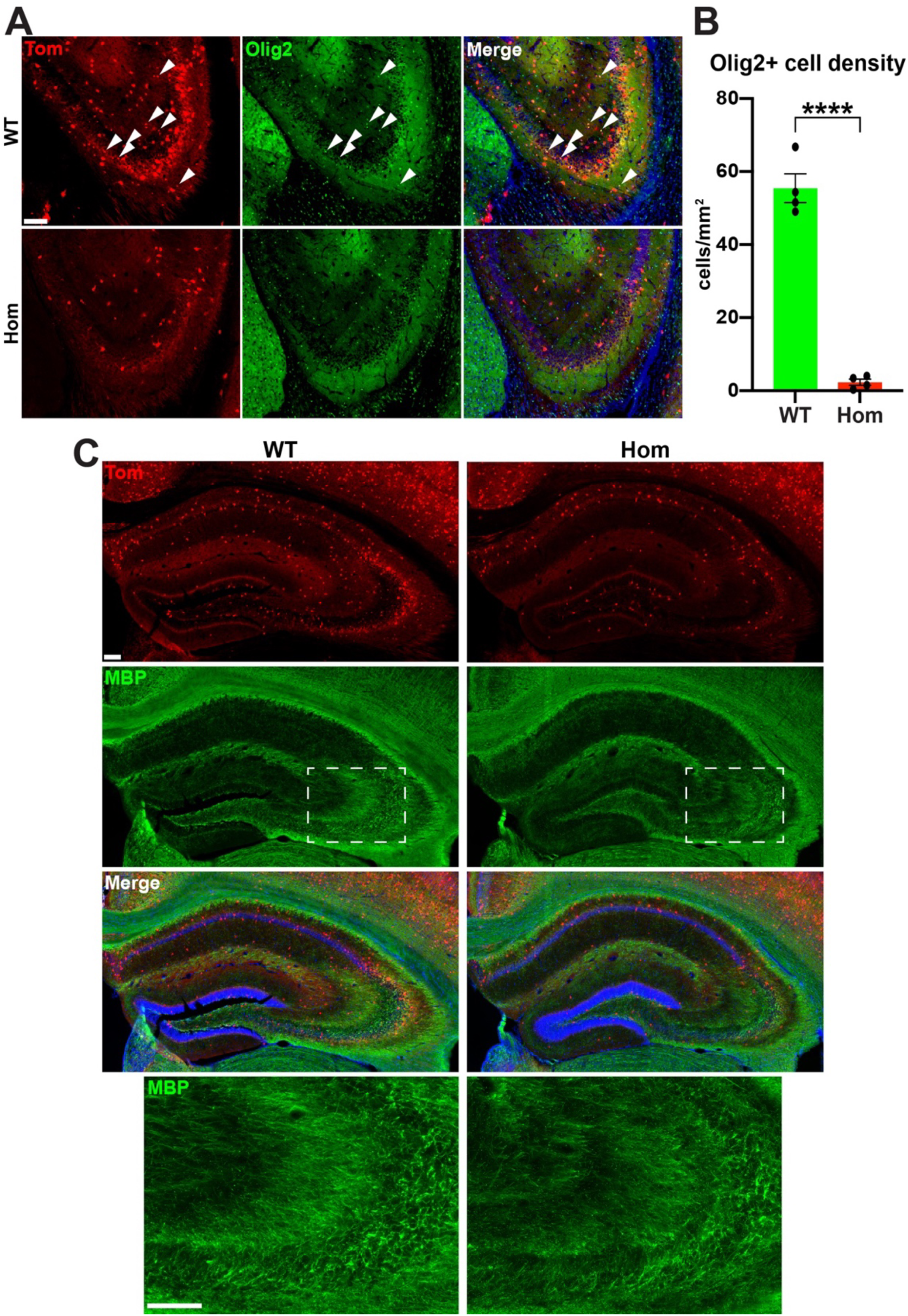
Loss of MGE-derived oligodendrocytes in H3.3K4M Hom mice. **A.** Representative images stained for Olig2 (green) through the CA3 showing loss of small cell body Tom+/Olig2+ MGE-derived oligodendrocytes (white arrowheads) in H3.3K4M Hom mice. **B.** Quantification of Tom+/Olig2+ oligodendrocytes in CA3 region of WT and Hom mice. Standard 2-tailed t-tests: ****p≤ 0.0001. **C.** Representative image showing reduction of myelin basic protein (MBP) in CA3 of H3.3K4M Hom mice. Scale bars = 50 µm.

**Figure S3.**
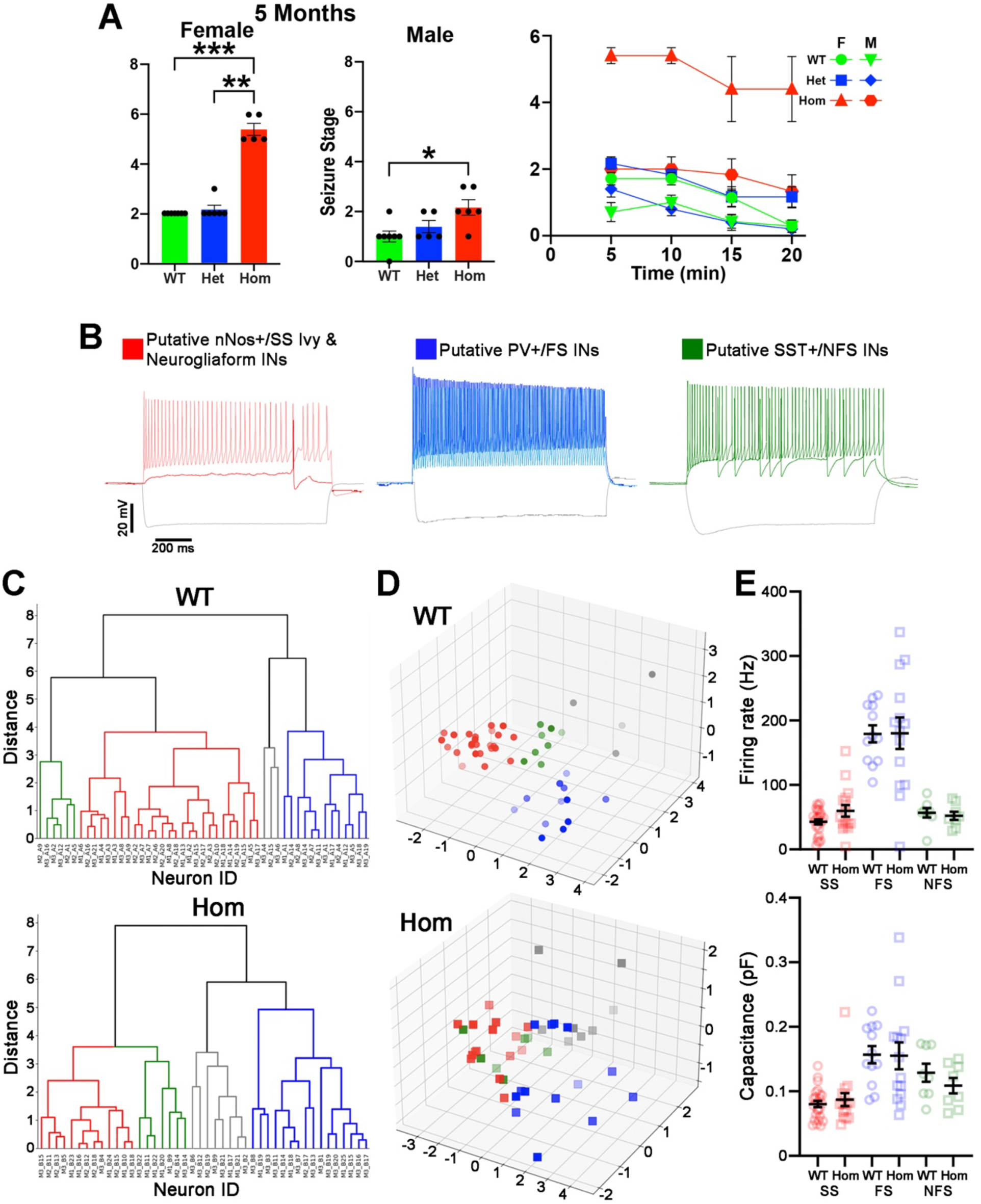
Altered intrinsic properties and network connectivity of hippocampal interneurons in H3.3K4M Hom mice. **A.** Maximum scores (seizure stage, left) for males and females following PTZ injection (20 mg/kg) at 5 months, and along the 20-minute observation period (right). **B.** Representative patch-clamp recording traces form putative nNos+/slow-spiking (SS, red), PV+/Fast Spiking (FS, blue) and SST+/non-FS (NFS, green) interneurons. **C-D.** Unbiased hierarchical clustering dendrograms (C) and PCA plots (D) of all hippocampal interneurons recorded from H3.3K4M WT (n = 49 cells) and Hom (n = 44 cells) mice. Cells that could not be classified are gray. **E.** Graphs depicting firing rate (top) and capacitance (bottom) for SS (red), FS (blue) and NFS (green) cells. Significant increase in variance in the FS population evaluated by F test in membrane capacitance (p = *) and firing rate (p = *) is observed in H3.3K4M Hom mice compared to WT. Kruskal-Wallis followed by Dunn’s multiple comparisons (A: left, middle) or two-way ANOVA followed by Tukey’s multiple comparison tests (A: right) were performed: * = p ≤ .05, ** = p ≤ .005, *** = p ≤ .0005.

**Figure S4.**
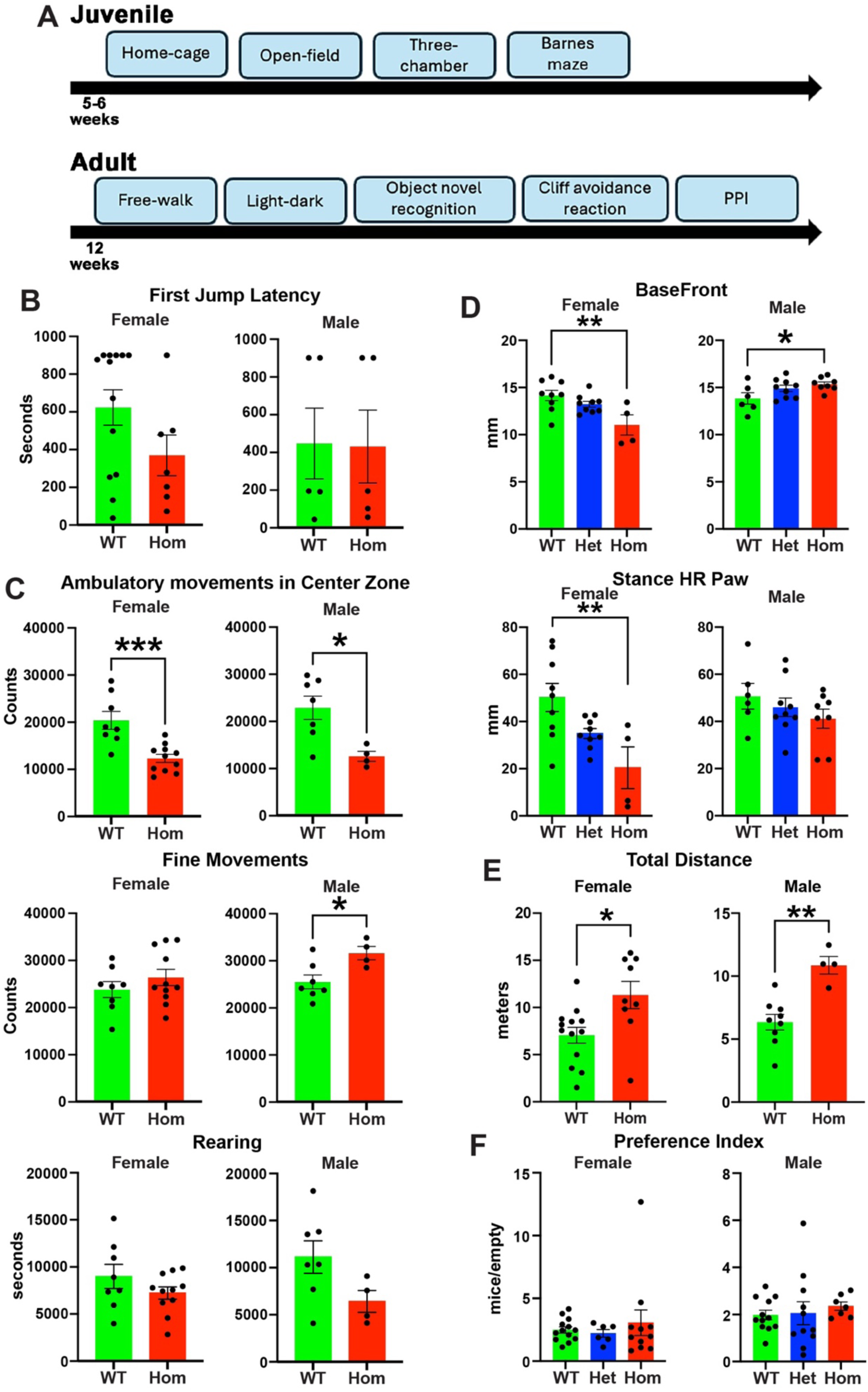
Increased anxiety and impaired locomotion in H3.3K4M Hom mice. **A.** Summary of behavior assays performed in juvenile and adult mice. **B.** First jump latency in the Cliff Avoidance Reaction Test. **C.** Beam break counts of ambulatory movements in the center zone (top), fine movements in the whole zone (middle) and rearing movements (bottom) over 4 days in the home cage test. **D.** BaseFront (top) and stance of hind right paw (bottom) in the free walk test. **E.** Total distance traveled by mice in the Barnes maze test. **F.** Preference index in the three-chamber test. All stats are one-way ANOVA followed by Tukey’s multiple comparison tests when WT, Het and Hom mice (D, F); standard 2-tailed t-test when only WT and Hom mice tested (B, C, E): * = p ≤ .05, ** = p ≤ .005, *** = p ≤ .0005.

**Figure S5.**
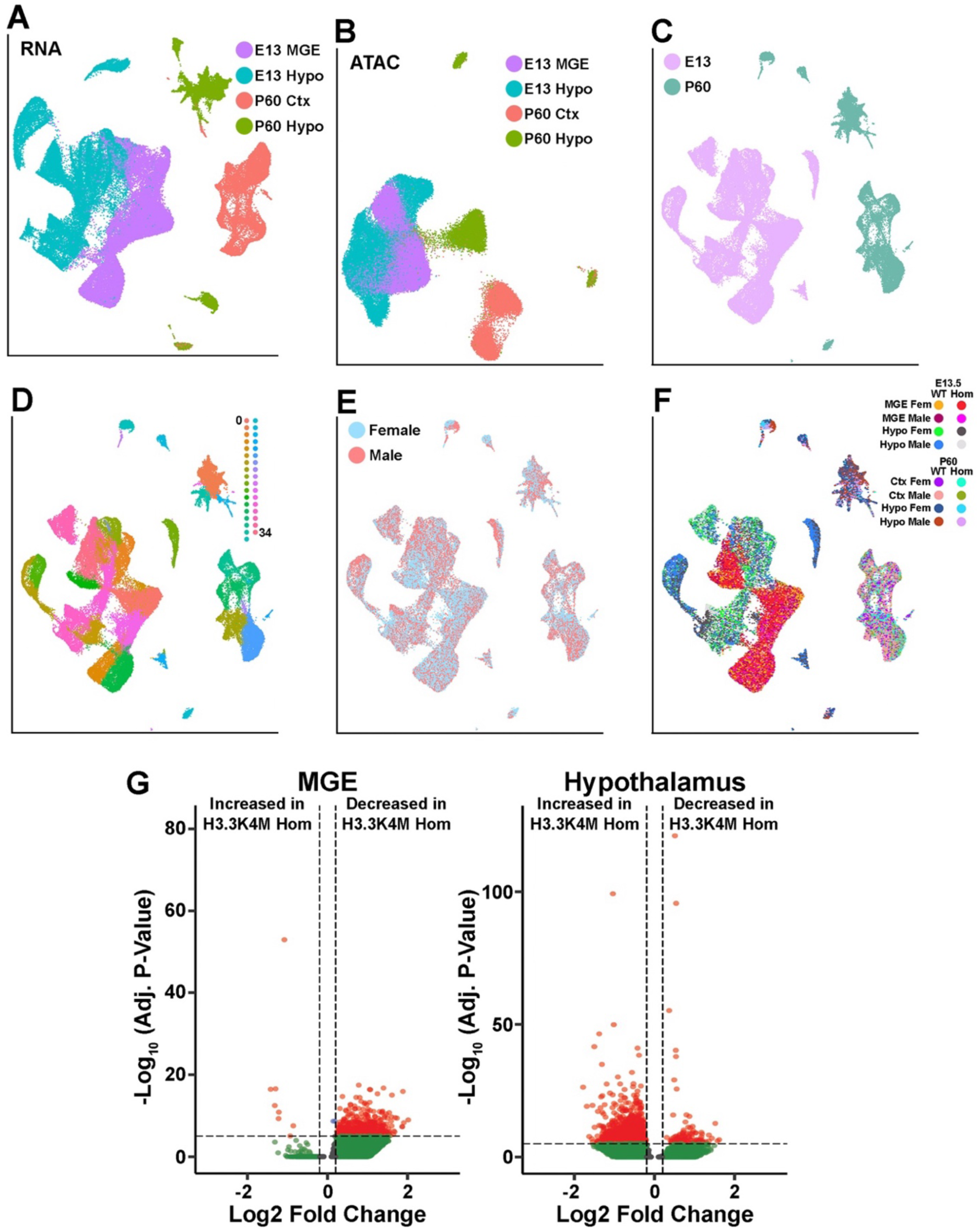
Total integrated Multiome-seq (snRNA-seq and snATAC-seq) data of 16 samples. **A-B.** UMAP plots of E13.5 MGE, E13.5 hypothalamus, P60 cortex and P60 hypothalamus annotated by age and tissue of the RNA-only (A) and ATAC-only (B). **C-F.** Integrated RNA and ATAC UMAP plots of E13.5 MGE, E13.5 hypothalamus, P60 cortex and P60 hypothalamus annotated by age (C), putative cell clusters (D), sex (E) and library id (F). **G.** Volcano plots depicting differentially accessible peaks from snATAC data in embryonic MGE (left) and hypothalamus (right) with combined sex.

**Figure S6.**
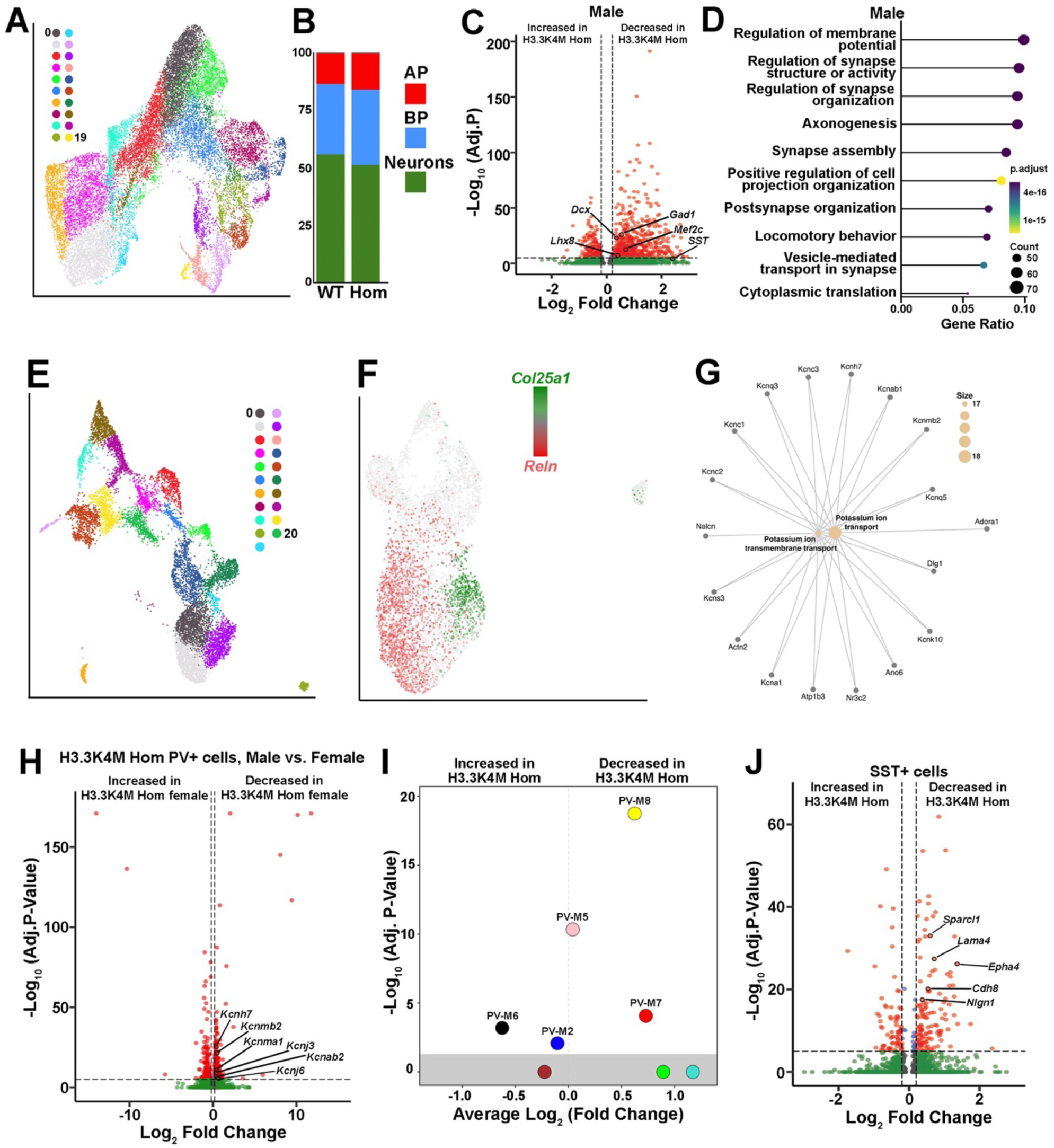
Altered transcriptome in MGE-derived interneurons associated with cell fate and seizure. **A.** UMAP plots of E13.5 MGE integrated single nuclei RNA- and ATAC-seq annotated by Seurat clusters. **B.** Proportions of E13.5 apical progenitors (AP), basal progenitors (BP) and neurons. **C.** Volcano plot depicting DEGs of male E13.5 MGE between H3.3K4M WT and Hom mice. **D.** clusterProfiler GO enrichment top biological processes for DEGs of male E13.5 MGE between H3.3K4M WT and Hom mice. **E.** Integrated UMAP plots of MGE-derived interneurons in P60 cortex annotated by Seurat clusters. **F.** UMAP plot of PV+ interneurons depicting *Reln* and *Col25a1* expression. **G.** Category netplot visualizing the GO terms of potassium ion transport and potassium ion transmembrane transport. **H.** Volcano plot depicting DEGs of P60 H3.3K4M Hom PV+ interneurons between female and male. **I.** Volcano plot depicting module eigengene differences identified by co-expression analysis in PV+ population between H3.3K4M WT and Hom mice. **J.** Volcano plot depicting DEGs of P60 SST+ interneurons between H3.3K4M WT and Hom mice.

**Figure S7.**
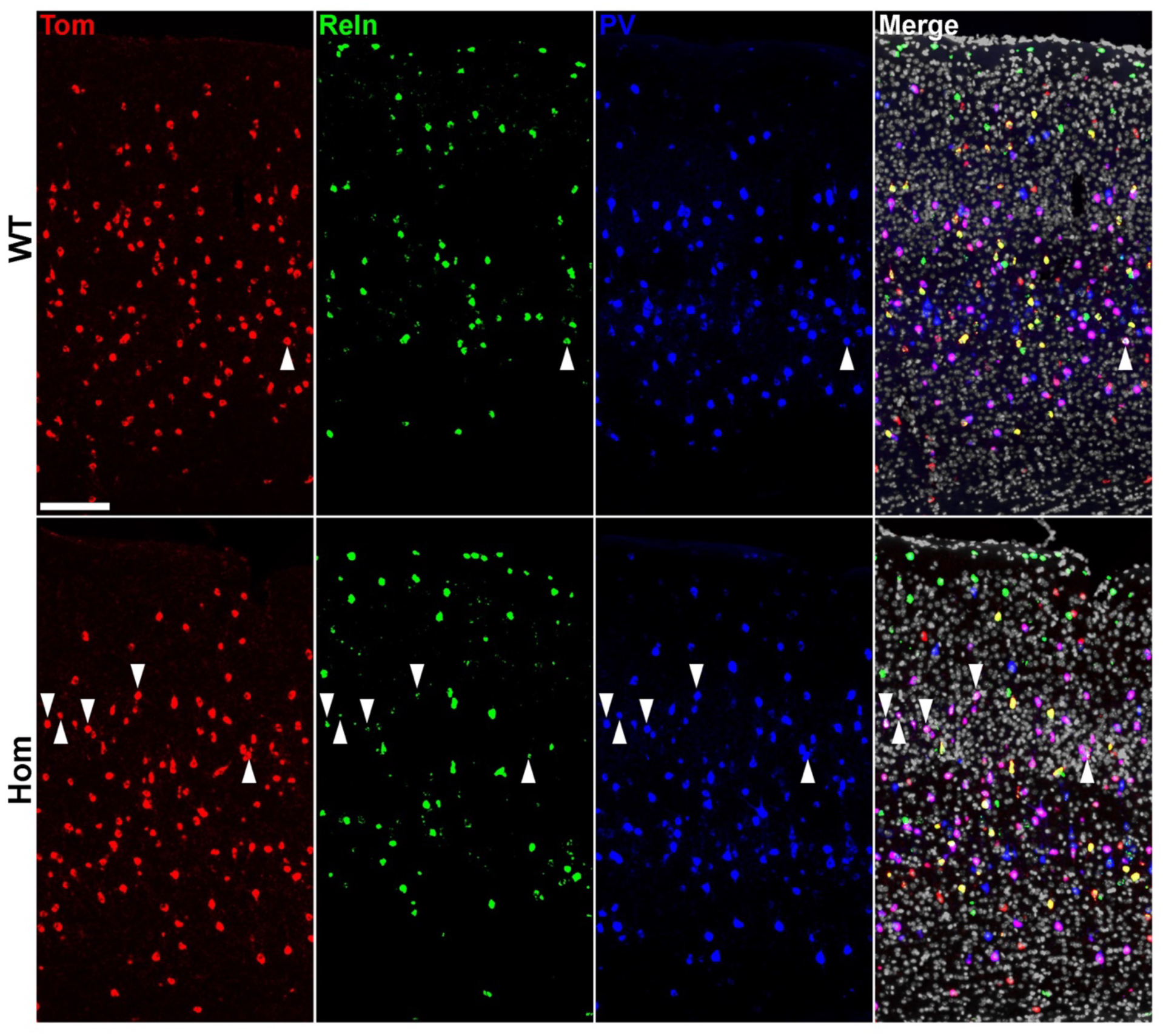
Increased PV+ / Reln+ in H3.3K4M Hom mice. Representative images of adult cortex showing increased number of Tom+/PV+/Reln+ cortical interneurons (white arrowheads) in H3.3K4M Hom mice compared to WT. Scale bar = 100 µm.

**Figure S8.**
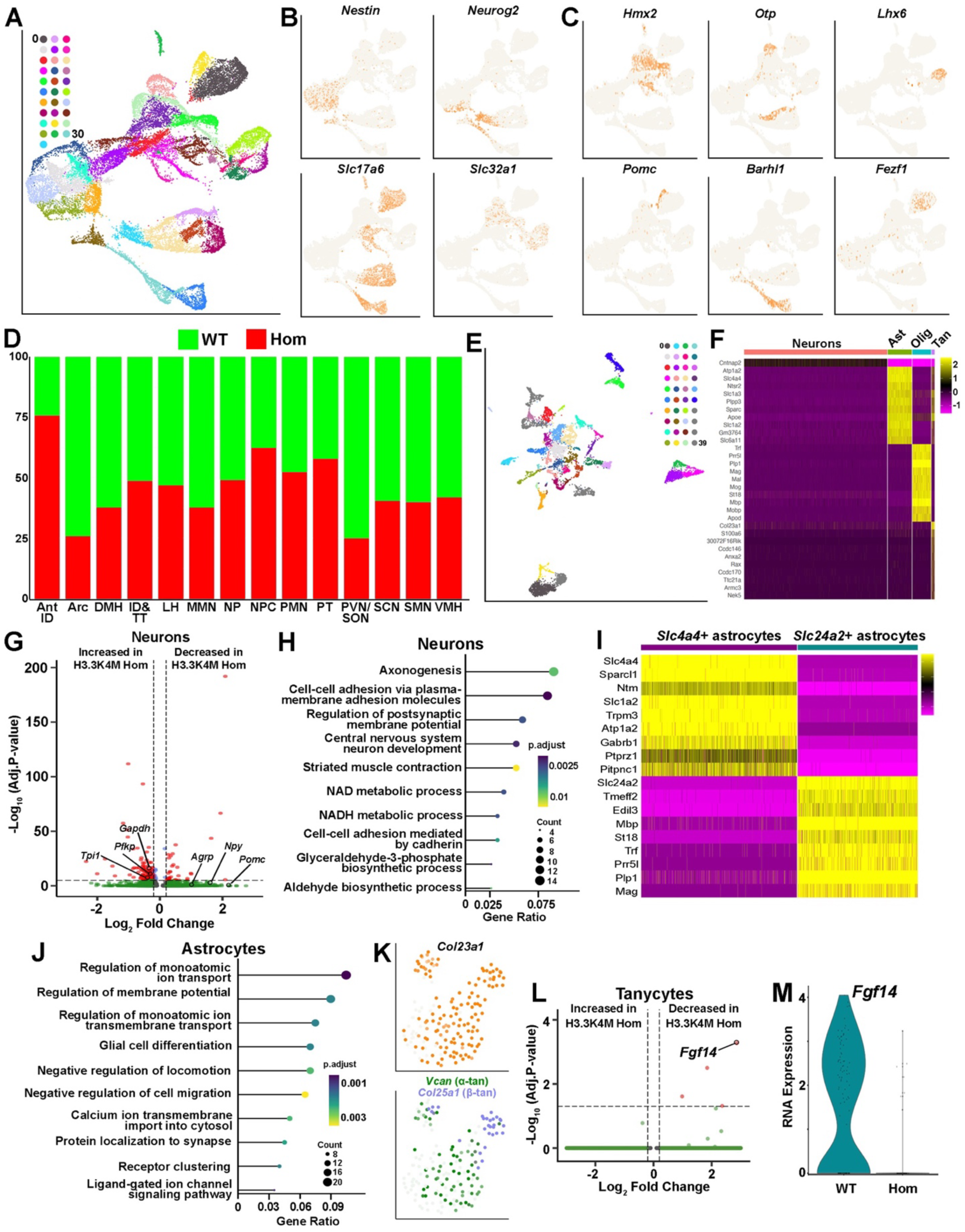
Transcriptome changes in numerous hypothalamic cell types in H3.3K4M Hom mice. **A.** Integrated UMAP plots of E13.5 hypothalamus annotated by Seurat clusters. **B.** Markers for apical progenitors (*Nestin*), neuronal precursors (*Neurog2*), postmitotic glutamatergic neurons (*Slc17a6*) and postmitotic GABAergic neurons (*Slc32a1*). **C.** Markers of region-specific hypothalamic nuclei PMN (*Hmx2*), DMH and PVN/SON (*Otp*), ID & TT (*Lhx6*), ARC (*Pomc*), SMN (*Barhl1*) and VMH (*Fezf1*). **D.** Relative proportions of hypothalamic nuclei/regions between H3.3K4M WT and Hom mice. **E.** Integrated RNA and ATAC UMAP plot of P60 hypothalamus annotated by Seurat clusters. **F.** Heatmap depicting critical genes that define astrocytes, oligodendrocytes and tanycytes. **G-H.** Volcano plot depicting DEGs of P60 hypothalamic astrocytes (G) and clusterProfiler GO enrichment of biological process for DEGs of P60 hypothalamic astrocytes (H). **I.** Heatmap depicting critical genes that define two astrocyte subtypes. **J.** clusterProfiler GO enrichment items of biological process for DEGs in P60 astrocytes. **K.** UMAP plot of all *Col23a1*+ tanycytes (top), and *Vcan*+ α-tanycytes (green) and *Col25a1*+ β-tanycytes (blue, bottom). **L.** Volcano plot depicting DEGs of P60 tanycytes. **M.** Violin plots showing significant downregulation of *Fgf14* in H3.3K4M Hom tanycytes. PMN, premammillary nucleus; VMH, ventromedial hypothalamus; LH, lateral hypothalamus; DMN, dorsomedial nucleus; Prethal, Prethalamus; ARC, Arcuate nucleus; ID&TT, intrahypothalamic diagonal & tuberomammillary terminal; SMN, supramammillary nucleus; PVN/SON, paraventricular nucleus/supraoptic nucleus; Ant ID, Anterior intrahypothalamic diagonal; SCN, suprachiasmatic nucleus; MMN, mammillary nucleus.

**Figure S9.**
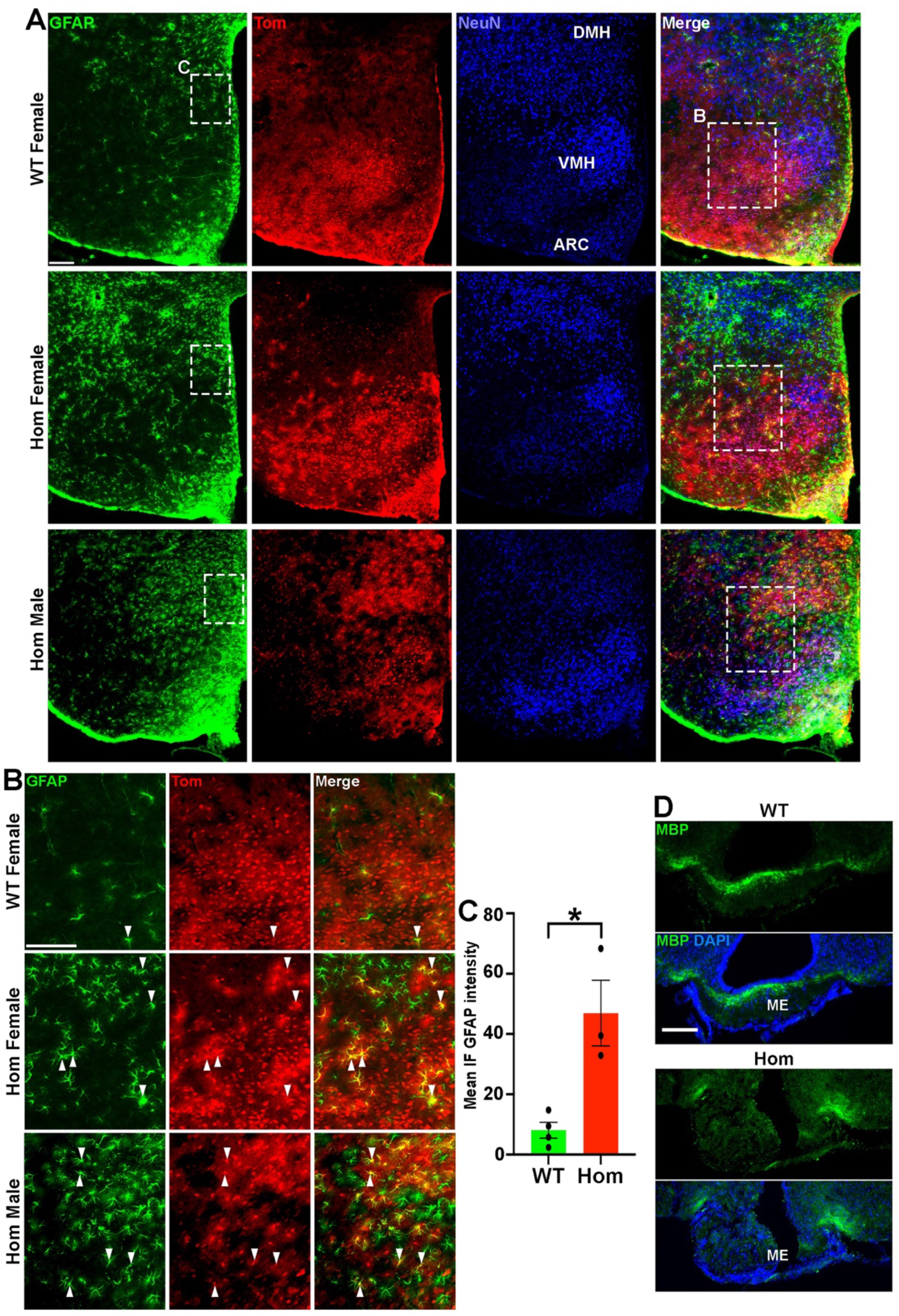
Dysregulation of glia cells in adult hypothalamus of H3.3K4M Hom mice. A-**C.** Representative images of adult hypothalamus from H3.3K4M WT and Hom male and female mice stained for GFAP (green), NeuN (blue), which Nkx2.1-linage cells expressing tdTomato. Scale bar = 100 μm. White rectangles in A indicate higher magnification regions (B), and regions used to measure immunofluorescence (IF) intensity of GFAP signal (C). **D.** Representative images of adult median eminence (ME) from H3.3K4M WT and Hom mice stained for MBP (green) and DAPI (blue). Scale bar = 100 μm. ARC, Arcuate nucleus; VMH, ventromedial hypothalamus; DMH, dorsomedial hypothalamus.

